# Characterization of the IGH locus and tissue specific immunoglobulin repertoires in turbot (*Scophthalmus maximus*)

**DOI:** 10.64898/2026.06.30.735498

**Authors:** Rocío Toucedo, Yixin Zhu, Sarai Moledo, Francisco Gambón, Pierre Boudinot, Ysabel Santos, Susana Magadán

**Affiliations:** Immunology Laboratory, CINBIO, University of Vigo, Campus Lagoas Marcosende, Vigo, Spain; Department of Computational Biology, Cornell University, Ithaca, NY, USA; Department of Quantitative and Computational Biology, University of Southern California, Los Angeles, CA, USA; Virologie et Immunologie Moleculaires, INRAE, Universite Paris, Saclay, Jouy-en-Josas, France; Department of Microbiology and Parasitology, IAQBUS, Universidade de Santiago de Compostela, 15782 Santiago de Compostela, Spain

**Keywords:** Turbot, IGH locus, immunoglobulin repertoire, 5′-RACE sequencing, B-cell clonotype diversity, skin immunity

## Abstract

Turbot (*Scophthalmus maximus*) is an important aquaculture species, but the genomic organization and expressed diversity of its antibody repertoire remain incompletely characterized. In this study, we annotated the immunoglobulin heavy chain (IGH) locus using the haplotype resolved fScoMax1.1 genome assembly, and we used this as a reference to profile the expressed turbot IgM, IgD and IgT repertoires in skin and spleen. The primary IGH locus was located on chromosome 19, spanned approximately 72 kb, and contained 25 IGHV genes, including 24 functional genes and one pseudogene, together with three IGHD, seven IGHJ and three IGHC genes corresponding to IgT, IgM and IgD. Comparison with the alternate fScoMax1.1 haplotype and a second turbot genome assembly showed conserved IGHD, IGHJ and IGHC content, whereas IGHV gene number differed among assemblies. High throughput 5RACE repertoire sequencing revealed isotype and tissue associated differences in expressed IGH diversity. IgM represented the dominant productive repertoire in both skin and spleen and showed the highest clonotypic diversity, particularly in spleen. IgD displayed an intermediate profile, whereas IgT was more enriched in skin and exhibited the strongest clonal restriction. IGHV subgroup usage was dominated by IGHV3 in IgM and IgD, whereas IgT showed a distinct profile characterized by preferential use of IGHV4, especially in skin. Gene level analysis further showed broad IGHV-IGHJ pairing in IgM and IgD, with preferential use IGHJ3 segment, while IgT sequences paired exclusively with IGHJT. Clonotype sharing between skin and spleen was isotype dependent, being strongest for IgT, intermediate for IgM, and negligible for IgD, suggesting that clonal expansion did not necessarily predict inter tissue trafficking. Together, these results provide a curated genomic and expressed repertoire framework for turbot IGH genes and reveal isotype specific organization of antibody diversity, with IgT displaying a particular repertoire pattern.

## Introduction

The adaptive immunity, a hallmark of vertebrate evolution, relies on the clonal expression of somatically diversifying antigen receptors on lymphocytes, the immunoglobulins (IG) or antibodies, in B cells, and T cell receptors (TR) in T cells. This process generates diversity and antigen specific recognition far beyond those offered by the allelic variation or alternative splicing processes. Antibodies are proteins composed of two heavy (H) chains and two light (L) chains that are encoded by the IGH locus and IGL locus, respectively. In most of the studied vertebrates, including teleost, the IGH loci have a translocon configuration. Which means a succession of IGHV, IGHD and IGHJ genes, followed by exons that code for the constant region (CH), arranged as (V)n-(D)–n -(J)n-(CH)n, where n represents the number of gene segments or exons (Lefranc and Lefranc, 2001). During the differentiation of B lymphocytes, the somatic recombination of one IGHV, one IGHD and one IGHJ leads to the assembly of an Ig gene encoding the complete V domain. Thus, the native structure of the IGH locus has a major effect on adaptive immunity, determining the range of gene segment choices available for the VDJ recombination process giving rise to the diversity of antigen-receptor sequences, the possible antibody classes and effector functions available in a host species (Alt et al., 1984; Hsu, 2009; Lewis, 1994). Genomic studies have revealed extensive plasticity in the organization of adaptive immune receptor loci across vertebrates (Bradshaw and Valenzano, 2020; Garzón-Ospina and Buitrago, 2020; Yasuike et al., 2010). Recent evidence further indicates substantial intra-species genomic variation, which adds an additional layer of complexity to shape the adaptive immune receptor repertoire (Rodriguez et al., 2023; Safonova et al., 2022; Watson et al., 2019). This variation includes differences in gene copy number, allelic diversity, and structural rearrangements within receptor loci, which can alter the set of germline-encoded V-D-J gene segments available for recombination. The comparative analysis of multiple genome assemblies from the same species will help to determine the extent and structure of this intra-species diversity, allowing them to distinguish conserved from variable locus features, and accurately link germline architecture to inter-individual differences in repertoire composition and adaptive immune responses.

Teleost represent the most diverse group of vertebrates, with around 30000 species. They have adapted to a wide range of ecological niches, making them an intriguing model for studying the evolution of immune responses. While the characterized teleost IGH loci show a configuration in translocon (Bengtén et al., 2006; Bradshaw and Valenzano, 2020; Gambón-Deza et al., 2010; Magadán-Mompó et al., 2011; Yasuike et al., 2010), differences in the number of V-D-J genes, arrangement and isotype availability can be found in teleost IGH loci, even in species closely related (Bradshaw and Valenzano, 2020; Magadan et al., 2019b) which can involve unknown consequences for antibody diversity, antigen recognition, and immune response against pathogens and/or vaccines. Furthermore, the immunoglobulin gene diversity among teleost species makes it challenge the extrapolation of insights gained from stablished fish models (e.g., zebrafish and salmonids) to evolutionary distant teleost species, particularly with respect to how effective infection-induced antibody responses are generated and how such responses can be induced by vaccination.

Three classes of antibodies: IgM, IgD, IgT, have been identified in teleost (reviewed in (Bilal et al., 2021; Fillatreau et al., 2013)). Interestingly, the DH and JH segments expressed by IgT are located between the IGHV gene segments and the DH and JH segments expressed by IgM or IgD (Danilova et al., 2005; Fillatreau et al., 2013; Flajnik, 2005; Hansen et al., 2005). Thus, the VDJ rearrangement results in the development of mutually exclusive B-cell lineages displaying surface IgT or IgM/D, which can express the same repertoire of VH genes, but not the same DH or JH, which could affect the specificity of antigenic recognition, and the differential role of B lymphocytes that express IgM/IgD or IgT. The IgT has been proposed as the functional equivalent of mammalian IgA in teleosts, with a main role in mucosal tissues (Gomez et al., 2013; Magadan et al., 2015; Zhang et al., 2010). For instance, recent studies performed in rainbow trout, to characterize skin adaptive immune response of fish against parasite and bacteria, revealed that *Ichthyophthirius multifiliis* and *Flavobacterium columnare* infection induced an increase of pathogen specific IgT titers in rainbow trout skin mucus, with a local proliferation of IgT expressing B cells, but no IgM or IgD specific titers were detected (Xu et al., 2013; Zhang et al., 2021). Thus, in the rainbow trout model, skin adaptive immune responses largely resemble those reported in the gills and gut of rainbow trout, in which IgT seems to have a main role (Gomez et al., 2013; Parra et al., 2016; Xu et al., 2016; Zhang et al., 2010). While IgM represents the main Ig in the plasma of teleosts and the main player in systemic immune responses, the role of IgM in mucosal responses should not be underestimated. Recent studies showed increased intestine and skin IgM + B cells in *Tenacibaculum maritimum* infected turbot (Faílde et al., 2014), and in skin of bath vaccinated flounder (*Paralichthys olivaceus*) against *Vibrio anguillarum,* the mRNA expression of IgM and secretory polymeric Ig receptors (pIgR) are increased (Sheng et al., 2019). Suggesting that differences among species, routes of vaccine delivery, and the nature of stimulation, immunization versus infection, may influence the relative predominance of IgT compared with IgM/IgD responses.

Turbot (*Scophthalmus maximus*) is a relevant marine flatfish species for aquaculture industry, widely cultivated in Europe and Asia. Intensive aquaculture practices expose turbot to various bacterial, viral, and parasitic pathogens, making disease prevention a critical concern for the industry. Understanding the turbot immune system is therefore essential for developing effective vaccination strategies and health management practices. While the lack of specific reagents have hampered the assessment of the role and dynamics of turbot immune components, considerable efforts have been made to increase turbot genomic (Brittain et al., 2025; Figueras et al., 2016) and transcriptomic resources to identify components involved in the immune response against bacterial fish pathogens (Gao et al., 2016; Robledo et al., 2014; Ronza et al., 2016). The development of high-throughput sequencing technologies (HTS) has enabled the generation of genome assemblies from multiple individuals within a species and, it has become an invaluable tool for studying the adaptive immune response, including B cells and antibodies, in different contexts of basic and applied immunology, such as in infectious processes and after vaccination (Bashford-Rogers et al., 2019; Lindner et al., 2015, 2012; Ramadoss and Robinson, 2020; Safonova et al., 2022). The analysis of IG repertoires by HTS has also started to be applied in important farmed fish species, reflecting a growing interest for an accurate and comprehensive description of immune repertoire modifications during responses to common pathogens and vaccines (Castro et al., 2013; Magadan et al., 2019a; Perdiguero et al., 2019). Previous studies have characterized the basic components of turbot humoral immunity, including the identification of IgM (Gao et al., 2014) and IgT (Tang et al., 2018) heavy chains, and the distribution of Ig-positive cells in lymphoid organs (Fournier-Betz et al., 2000). Recent single-cell RNA sequencing has provided a comprehensive immune cell landscape of turbot, identifying distinct B cell populations in systemic and mucosal tissues, as well as a preliminary analysis of the adaptive immune receptor encoding loci (Chen et al., 2022). However, a comprehensive analysis of immunoglobulin repertoire diversity across tissues and isotypes has not been performed. In this study, we examined the genomic sequence of the IGH locus using the haplotype resolved fScoMax1.1 assembly and the ASM2237912v1 genome assembly. Comparative analysis revealed conserved IGHD, IGHJ and IGHC content across assemblies, whereas IGHV gene number varied among the three annotated IGH loci. This comprehensive and coherent IGH locus annotation supports the analysis of high-throughput repertoire sequencing data. In parallel, we characterized the expressed immunoglobulin heavy-chain repertoire of turbot across two tissues representing systemic and mucosal immune compartments, spleen and skin, respectively, and across the three major teleost Ig isotypes: IgM, IgD, and IgT. High-throughput 5′-RACE repertoire sequencing revealed clear isotype- and tissue-associated differences in expressed IGH diversity. IgM dominated the productive repertoire in both skin and spleen and displayed the highest clonotypic diversity, particularly in spleen. IgD showed an intermediate profile, whereas IgT was enriched in skin and exhibited the strongest clonal restriction.

## Material and Methods

### Annotation of IGH locus

Turbot genome assemblies (fScoMax 1.1; ASM2237912v1) were accessed through the Vertebrate Genome Project (https://vertebrategenomesproject.org/) or NCBI website (Brittain et al., 2025). In the case of fScoMax1.1, we analyzed the main assembly (GCA_963854745.1) and the alternate haplotype (GCA_963854755.1). To identify the chromosomes that embrace the IGH locus, previously published turbot IGH sequences (Gao et al., 2014; Tang et al., 2018) were used for BLASTn and tBLASTn searches. In fScoMax 1.1, the regions encoding for IGHC were located on chr. SUPER19 (OY978270.1) and scaffolds atg000433l_1 and atg002984l_1. While in ASM2237912v1 turbot genome assembly, the IGH locus is located on Chr17 (NC_061531.1). These chromosomes and scaffolds were selected for in depth analysis. IGH genes were searched using the BLAST tool at Galaxy website (https://usegalaxy.org) and visual analysis using SnapGene software (available at snapgene.com). The annotation and functionality of IGH genes were established according to IMGT (www.imgt.org) standards (Lefranc and Lefranc, 2001; Lefranc, 2014). To annotate IGH variable V, joining J, and diversity D genes, the recombination signal (RS) sequences were identified, respectively V-RS, J-RS, and 5’D-RS and 3’D-RS. Splice signals were used to determine the limits of the coding nucleotide sequences for the V (L- PART1 donor splice and L-PART2 acceptor splice) and the J (J-REGION donor splice). The following parameters were considered to determine the gene functionality: a) presence of appropriate splice sites, b) presence of RS sequence compatible with effective rearrangement, c) open reading frames which included conserved cysteine (CYS) and tryptophan (TRP) codons at positions 23 (1st-CYS), 41 (CONSERVED-TRP) and 104 (2nd-CYS) of IGHV genes. Briefly, a germline entity (IGHV or IGHJ gene) was considered functional (F) if the coding region has an open reading frame, no defect in splicing site or in RS, and presents key conserved amino acids; as open reading frame (ORF) if the coding region has an open reading frame with defects in the splicing sites or RS sequences, and/or changes of conserved amino acids that lead to incorrect folding; or annotated as pseudogene (P) if the coding region has stop codon or frameshift mutations. Chromosomal coordinates given in the turbot IGH annotation file (**Supplementary file 1**) include the recombination signal (RS) sequence and coding regions of IGH genes.

Turbot IG genes were named following similar principles as in our recent work on salmonid IGH, TRA/TRAD and TRB loci (Boudinot et al., 2023; Edholm et al., 2021; Magadan et al., 2019b). Specifically, the IGHV genes were assigned to subgroups using a threshold of 75% nucleotide identity. The variation in the number and position of IGHV genes among the three studied assemblies (Figure 1) prevented us from proposing a positional nomenclature. Thus, each IGHV gene is named according to its subgroup and its sequential order of identification within that subgroup. For example, IGHV3-2 denotes the second annotated IGHV gene assigned to subgroup 3 (Figure 1 and Supplementary File 2). The annotation of turbot IGH locus and proposed nomenclature were submitted to the Immunoglobulins (IG), T cell receptors (TR), and major histocompatibility (MH) Nomenclature Sub-Committee of the International Union of Immunological Societies (IUIS) Nomenclature Committee (https://iuis.org/committees/nom/nomenclature-sub-committees/t-cell-receptor-and-immunoglobulin-nomenclature-sc/) for review and formal acceptance.

**Figure 1.**
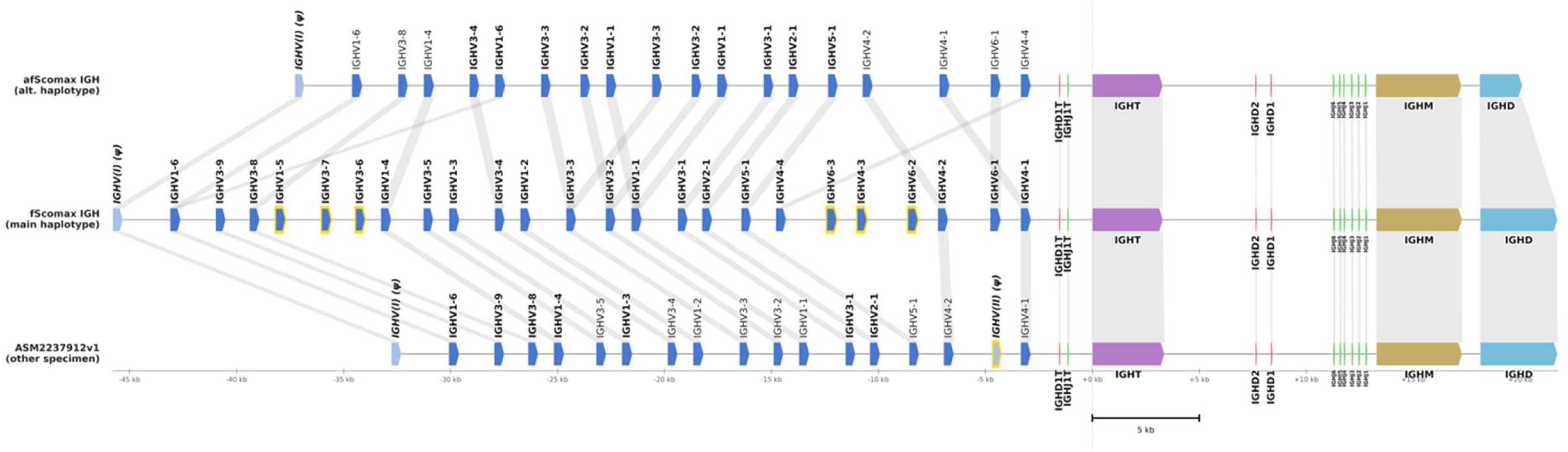
Organization of the IGH locus in turbot (*Scophthalmus maximus*) across three genome assemblies. The IGH locus is shown for the fScoMax1.1 primary haplotype (GCA_963854745.1; top), the fScoMax1.1 alternative haplotype (GCA_963854755.1; middle), and the ASM2237912v1 assembly (GCA_022379125.1; bottom). All three loci are aligned at the start of the IGHT constant region gene (dashed vertical line, position 0), with genomic coordinates in kilobases (kb) relative to this anchor. Gene segments are directional arrows colour-coded by type: IGHV (blue), IGHD diversity segments (red), IGHJ joining segments (green), IGHT (purple), IGHM (gold), and IGHD constant region (light blue). Pseudogenes are shown in light blue and labelled with the ψ symbol. Gene segments present in only one assembly are highlighted with a yellow border. Grey shaded linkers connect gene segments between adjacent assemblies. Scale bar, 5 kb.

CloseRead (Zhu et al., 2025) analysis was performed on the turbot IGH region from the fScoMax1.1 (both primary and alternative haplotype assemblies) and ASM2237912v1 genome assemblies. CloseRead was run with default parameters using user-provided gene annotation files to evaluate and compare IGH locus assembly quality across the two assemblies.

### Phylogenetic analysis

Phylogenetic analysis was performed on IGHV sequences. Sequences were aligned using ClustalW in the MEGA-7 software (Kumar et al., 2016). The phylogenetic trees were then constructed using the Neighbor-joining method in MEGA 7, and bootstrapped 2000 times.

### Samples and RNA extraction

Spleen and skin were collected from four non vaccinated turbot (50-75g). Tissues were aseptically harvested, kept in RNA Later at -20°C until used. Total RNA was individually prepared from spleen and skin by disruption of the tissues in Trizol reagent (Life Technologies). Then, RNA was purified and DNA removed using RNeasy Mini-Plus kit (Qiagen). RNA concentration was determined by Nanodrop measure and stored at -80°C until further use.

Non vaccinated turbot were reared at 16±1°C at the Aquarium of the Faculty of Biology of the University of Santiago de Compostela. All experiments were carried out in accordance with the European Union guidelines for the handling and welfare of laboratory animals (https://ec.europa.eu/environment/chemicals/lab_animals/index_en.htm). The experimental protocols were approved by the USC institutional ethics committee (15004/2022/004). Fish were sacrificed by overexposure to MS-222 (Tricaine) followed by destruction of the brain. Tissue samples of the spleen and skin (approximately 5 cm^2^) were taken from each fish and stored in RNA later (Thermofisher) at -80°C until RNA extraction.

### MiSeq library preparation and sequencing

Libraries were generated following previously described 5′-RACE-based approaches for immunoglobulin repertoire sequencing (Al Hayek et al., 2025; Turchaninova et al., 2016). One microgram of total RNA from the spleen or skin was used as starting material for cDNA synthesis. All primers for this study can be found in **Supplementary File 2.** Reverse transcription (RT) was carried out using isotype specific primers targeting the Cµ2, Cδ2, or Cτ1 constant region domains, together with SMARTer technology, which introduces 5′ poly-cytidine overhangs upstream of the untranslated region. A template-switch oligonucleotide, Rd2p_UID_TS, containing a 15-nucleotide random sequence used as a unique molecular identifier (UMI) and a sequence complementary to the Illumina adaptor, was then incorporated.

The resulting cDNA was purified using Mag-MAX^TM^ Pure Bind Beads (Applied Biosystems) at a ratio of 1:1.2 (sample:beads) following the manufacturer’s instructions and eluting the cDNA in 30 μl of H_2_0. The purified cDNA was further amplified in a two-stage PCR in 50 μl volume reaction using with 5 PRIME HotMaster Taq DNA Polymerase (Quantabio). The first PCR was performed using an isotype-specific primer complementary to the Cµ1, Cδ2, or Cτ1 constant-region domain, together with a primer complementary to the Illumina adaptor sequence (**Supplementary File 2**). Cycling conditions were as follows: initial denaturation at 94°C for 3 min; 25 cycles of 94°C for 30 s, 59°C for 30 s, and 72°C for 30 s; followed by a final extension at 72°C for 10 min. PCR products were loaded onto a 1% agarose gel. After electrophoresis, the IGHV-Cmu or IGHV-Ctau products at 450–600 bp, and IGHV-Cdelta products at 700 bp were purified by double sized selection process using Mag-MAX^TM^ Pure Bind Beads (Applied Biosystems) following manufacturer recommendations and eluted in 30 μl of H_2_0. A second PCR reaction was then performed to add the remaining Illumina adaptor sequences and unique sample indexes, as previously described (Magadan et al., 2018), using the primers listed in **Supplementary File 2.** Cycling conditions were as follows: initial denaturation at 94°C for 3 min; 15 cycles of 94°C for 30 s, 64°C for 30 s, and 72°C for 30 s; followed by a final extension at 72°C for 10 min. The second PCR products from the same first PCR were pooled and purified using Mag-Bind beads at a ratio of 1:1 and eluted in 20 μl of H_2_0. An aliquot of the purified samples was run on a 1% agarose gel to ensure amplification of the correct size (∼600 bp). The PCR product concentration in each library was determined using a Qubit fluorimeter (Invitrogen) and the quality of pooled library was checked using an Agilent 2100 Bioanalyzer (Agilent Technologies). Sequencing was performed at the Sequencing Service of the University of Salamanca (Spain) on the Illumina MiSeq instrument using paired-end 2 x 300 bp runs and the MiSeq Reagent Kit v3 (600 cycles) (Illumina).

### Raw data processing and IG repertoire analysis

Initial data analysis involved the adaptor trimming performed Cutadapt 3.2. Reads were filtered and merged to create consensus sequences using pRESTO on Galaxy (Galaxy Community, 2024; Vander Heiden et al., 2014). First, raw reads with a Phred score lower than 20 were filtered out and primers were masked. Reads where the primers could not be identified or had a poor alignment score were removed. Next, paired-end reads were aligned and merged (maximum error of Assemble Pairs align set to 0.02). The consensus sequences were annotated using IgBLAST v1.22.0 (Ye et al., 2013) with a custom turbot IG germline database, which included the 24 functional IGHV genes, 3 IGHD genes, and 7 IGHJ genes annotated in the primary haplotype fScoMax1.1 assembly. The Framework region (FR) and complementarity-determining region (CDR) anchor points were defined following the standardized IMGT numbering positions (Lefranc and Lefranc, 2001). This information was used to create an IgBLAST compatible database. IgBLAST was configured to output results in AIRR (Adaptive Immune Receptor Repertoire) format, which provides standardized fields for IGHV, D, and J gene assignments, alignment coordinates, and productivity assessment (Eugster et al., 2022).

IGHV–IGHJ pairing analysis. For each isotype (IgM, IgD, IgT) and tissue (skin, spleen), productive sequences with unambiguous single V gene and J gene assignments were pooled across the four specimens (C3, C5, C7, C8). IGHV–IGHJ counts were aggregated and visualized as heatmaps with colour intensity representing log₁₀(count + 1) using the cividis colour scale. Analyses were performed in Python using pandas and matplotlib.

For further analysis, sequences sharing the same IGHV gene, IGHJ gene, and CDR3 nucleotide sequences were grouped into the same clonotype. Sequences without IGHV or IGHJ alignment, with internal stop codons or CDR3 associated junction without conserved cysteine (which marks the 3’end of FR3) and tryptophane (which marks the 5’of FR4) were excluded from downstream clonotype analysis. Rarefaction curves were generated using the vegan R package to assess sampling completeness, with 100 random subsampling iterations at each depth. Diversity metrics were calculated after rarefaction to the minimum sample size within each tissue (Skin: 879 sequences; Spleen: 4,135 sequences) to enable fair comparison across samples with different sequencing depths. Richness (observed number of clonotypes), Shannon diversity index, Simpson diversity index, Gini coefficient, and Chao1 estimator were calculated for each sample. Shared clonotypes between tissues were identified by matching IGHV-IGHJ-CDR3 combinations within each fish. Analyses were performed in R using vegan, ggplot2, and ComplexHeatmap packages.

## Results

### Turbot IGH locus

The newly released primary turbot haplotype genome assembly (fScoMax1.1, GCA_963854745.1) was used as a reference for the annotation of turbot IGH locus. This assembly is based on 41xPacBio and Arima2 Hi-C sequencing technologies. IGH genes were identified within a single locus on chromosome 19 (OY978270.1) in a forward (FWD) orientation. The turbot IGH locus spans approximately 72 Kb and comprises a total of 38 genes (**Figure 1**), including 25 IGHVs (24 functional and 1 pseudogen), 3 IGHDs, 7 IGHJ and 3 IGHC genes (IGHT, IGHM and IGHD). Comparison with the alternate haplotype assembly of fScoMax1.1 (GCA_963854755.1) and with a second primary turbot genome assembly, ASM2237912v (GCA_022379125.1), showed the same IGHC, IGHD and IGHJ gene content across assemblies (**Figure 1**). In all three assemblies, one IGHD gene of 13 nucleotides, was located upstream of IGHT gene, whereas two IGHD genes of 19 and 12 nucleotides were located upstream of IGHM-IGHD cluster (**Figure 1** and **Supplementary File 1**). Sequence analysis showed that the three IGHD genes present a G-rich stretch and, while IGHD1 and IGHD2 genes can be productively read in all coding frames, the IGHD1T can use five of the six coding frames (**Supplementary File 3**). In addition, the 5’D-RS and 3’D-RS sequences flanking the IGHD were conserved in all three IGHD genes, specifically the D-nonamers and the 3’D-heptamer. A total of seven IGHJ genes were identified in turbot IGH locus (**Supplementary File 2 and Suppl. File 3**), one linked to the IGHT locus and 6 belonging to the IGHM-IGHD cluster, all of them in forward transcriptional orientation. The length of turbot IGHJ genes ranged from 45 to 57 nucleotides. Among them, IGHJ3 and IGHJT were the shortest, comprising 45 and 47 nucleotides, respectively, whereas IGHJ1 was the longest, with a length of 57 nucleotides (**Supplementary File 2 and Suppl. File 3)**. All IGHJ genes showed strong conservation of sequence motifs as in other vertebrates including the amino acid Tryptophan-Glycine-X-Glycine (WGXG) J-MOTIF present in nearly all IGH J-REGION. Overall, turbot IGHJ genes displayed conserved recombination signal (RS) sequences, except for IGHJ6, whose nonamer sequence (5′-ACTATCTGT-3′) diverged from the consensus sequence 5′-GGTTTTTGT-3′. In contrast to the conserved IGHD-IGHJ-IGHC cluster, the number of annotated IGHV genes differed among the three turbot genome assemblies. While 25 IGHV genes were identified in the reference assembly fScoMax1.1, the analysis of the alternate haplotype of the same specimen and the second primary turbot genome assembly (ASM2237912v) resulted in 19 and 18 IGHV genes, respectively **(Figure 1).** The differences observed may reflect either genome assembly artefacts or true structural variation in the IGH locus among different specimens, as previously described in other vertebrates, including human (Rodriguez et al., 2023).

The discrepancies in the number of IGHV genes posed us to conduct an analysis with CloseRead (Zhu et al., 2025) to determine whether these assemblies accurately represent turbot IGHV gene cluster and their organization in the genome. As shown in **Figure 2**, across the primary and alternate fScoMax1.1-IGH assemblies, read depth was broadly consistent and mapping quality was strongly enriched for high-confidence alignments (MAPQ ∼60), supporting good contiguity and read concordance, a similar pattern was observed for ASM2237912v1 **(Supplementary File 4).** Importantly, fScoMax1.1 is haplotype-resolved and thus can preserve more allelic variation and enable identification of additional IGHV gene variants, whereas ASM2237912v1 represents a consensus assembly in which divergent haplotypic sequence may be partially collapsed. At the gene level, CloseRead indicated strong read support in both assemblies: 97% of genes in fScoMax1.1 showed 100% accuracy, and all genes in ASM2237912v1 exceeded 98% accuracy. A localized exception occurred near the fScoMax1.1 IGH 3′ J–constant region on SUPER_19 and at the terminus of atg000433l_1, where CloseRead flagged an increase in secondary alignments and a coverage drop respectively, consistent with increased local sequence complexity. Notably, because the alternate haplotype IGH locus is shorter than the primary, this signal is consistent with missing sequence in the alternate assembly within this interval.

**Figure 2.**
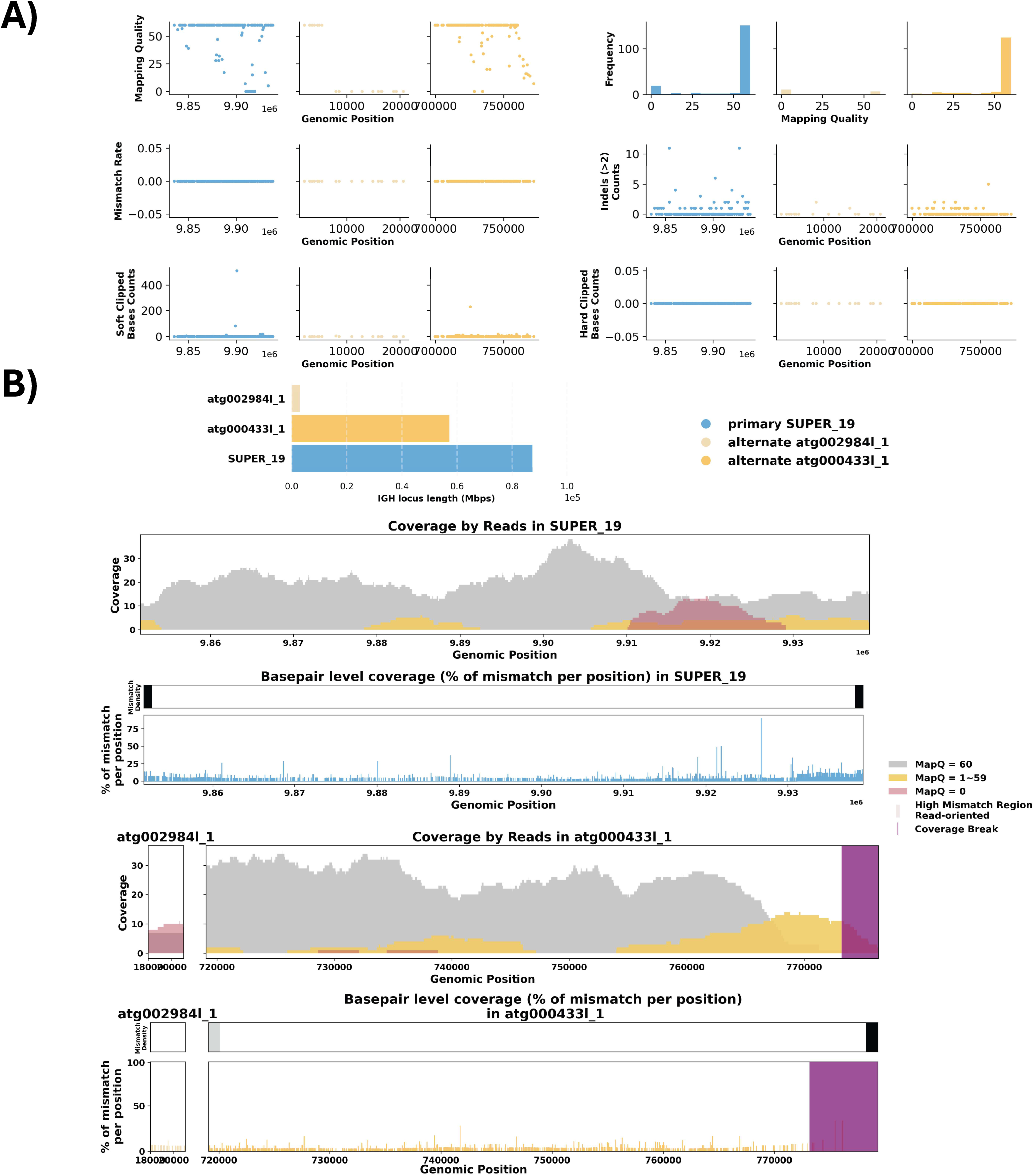
CloseRead results of IGH locus assembly in primary (IGH: SUPER 19) and alternate (atg002984I_1 and atg000433I_1) fScoMax 1.1 genome assemblies. A) Summary statistics of the read alignment, showing mapping quality across IGH loci for both haplotypes, with blue representing the primary assembly and yellow the alternate. Mismatch rates of reads, Number of reads with indels of consecutive length of at least 2 bp, and count of soft clipped bases in each read, B) IGH locus length in both the primary and alternate assemblies are compared using bar charts. The detailed analysis of alignment mismatch in both haplotypes includes (i) read coverage across the entire IGH loci, color-coded by mapping quality, and (ii) basepair-oriented mismatch rate, a heatmap above indicating the frequency of high mismatch rate base pairs.

### IGHV genes

We used the annotation of the IGH locus in the main fScoMax1.1 haplotype as a reference, as it contains the highest number of annotated IGHV genes **(Figure 1 and Supplementary File 2).** Based on nucleotide sequence identity across the V-REGION (75% threshold), the identified functional IGHV genes can be classified into **6** IGHV subgroups **(Figure 3)**. These subgroups vary in size, ranging from a single functional member (IGHV2 and IGHV5 subgroups) to 9 genes in IGHV3, 5 in IGHV1, 4 in IGHV4, and 3 in IGHV6 **(Figure 3)**. The three IGHV genes assigned to subgroup 6 exhibit 100% sequence identity, including the L1-INTRON region, which may indicate a potential assembly artifact. In addition, comparison of IGHV genes identified across the three analyzed turbot genome assemblies (main and alternate fScoMax1.1, ASM2237912v1) showed that all annotated functional IGHV genes can be consistently clustered into these six IGHV subgroups (**Figure 4**).

**Figure 3.**
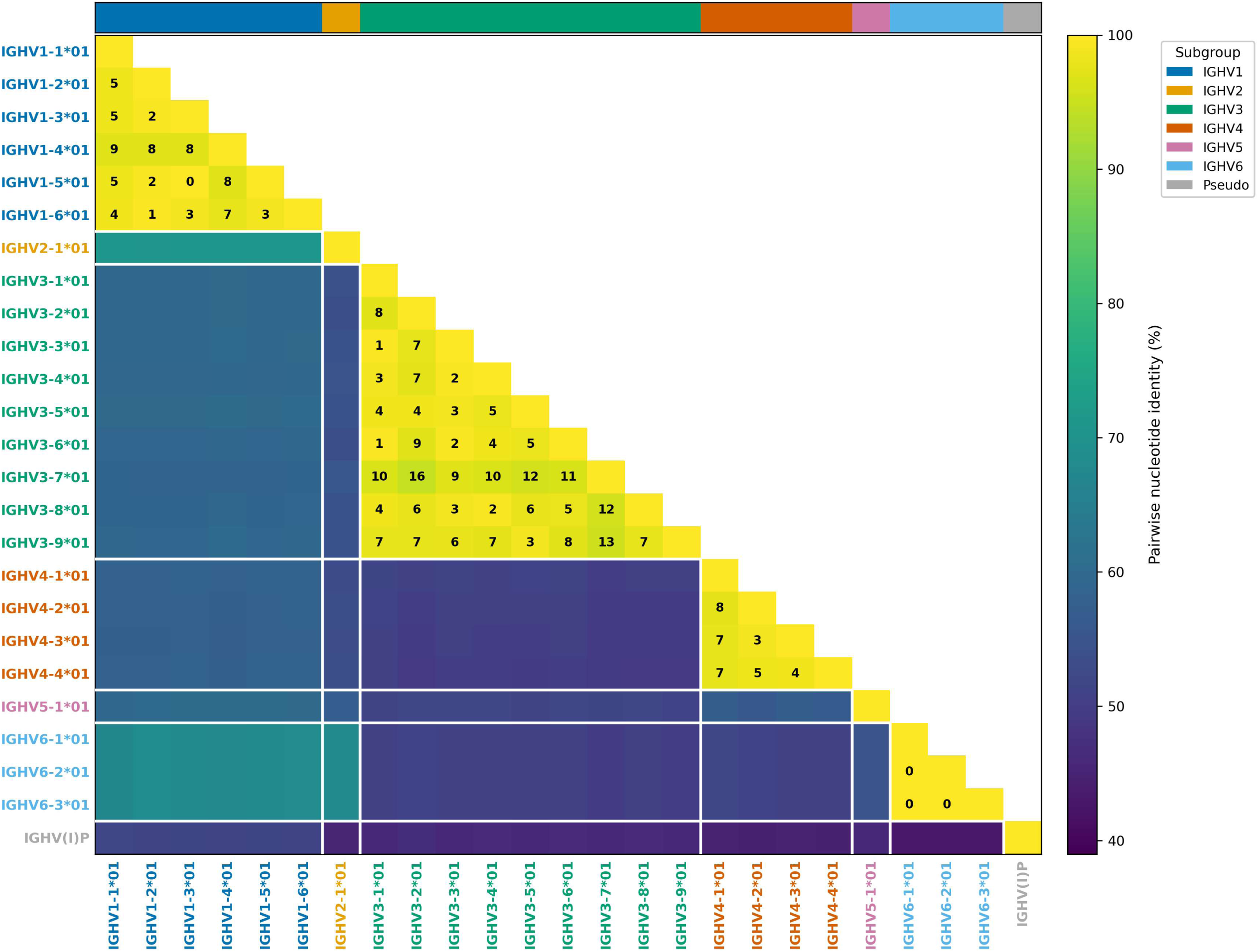
Pairwise nucleotide identity matrix of *Scophthalmus maximus* IGHV genes. The matrix displays the percentage of pairwise nucleotide identity between the 25 annotated IGHV genes in the main fScomax 1.1 haplotype assembly (GCA_963854745.1). The percentage of pairwise nucleotide identity was calculated from the multiple sequence alignment and excluding gap positions. Cells are coloured according to the viridis scale (dark purple = low identity, yellow = high identity). Numbers within cells indicate the number of differing nucleotides for gene pairs sharing >75% identity. Genes are ordered and grouped by subgroup (IGHV1–IGHV6), and the colour strip at the top indicates subgroup membership: IGHV1 in dark blue, IGHV2 in amber, IGHV3 in green, IGHV4 in vermillion, IGHV5 in Pink, and IGHV6 in sky blue. Pseudogenes are indicated in grey.

**Figure 4.**
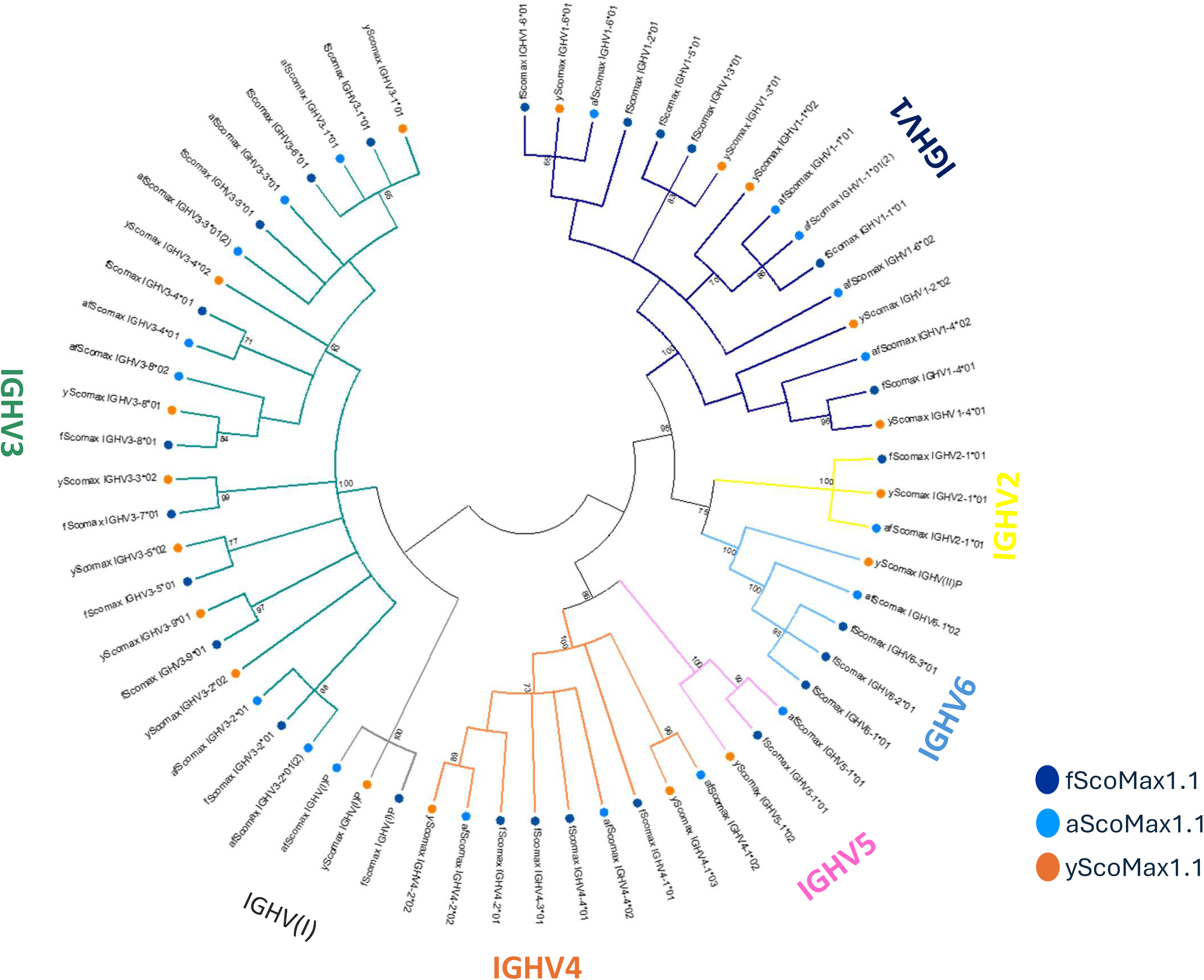
Phylogenetic analysis of turbot IGHV gene sequences. A phylogenetic analysis of the V-REGION of annotated turbot IGHV genes in the three genome assemblies (fScoMax1.1 primary haplotype (fScomax), the fScoMax1.1 alternative haplotype (afScomax), and the ASM2237912v1 assembly (yScomax)) was performed using nucleotide sequences, the Neighbor joining method and a boostrap based on 2000 replicates. A total of 62 IGHV sequences, 25 from fScomax, 19 from aScomax and 18 from yScomax were compared. Nodes with a boostrap support higher than 60% are indicated. The subgroup branches are represented in different colours.

We further characterized if the IGHV genes annotated in the alternate fScoMax1.1 and ASM2237912v1 IGH assemblies could correspond to new genes. We initially assumed that IGHV (V exon) nucleotide sequences assigned to the same gene should generally differ by no more than 2,5%. We performed a complete similarity matrix based on pairwise alignment of all pairs of sequences, and a clustering based on this criterion, and found a counterpart for 19 out 26 genes annotated in the main fScoMax1.1 assembly (**Figure 1 and Supplementary File 1**). Interestingly, members of the IGHV6 subgroup were only identified in fScoMax1.1 genome, being present in both the primary and alternate haplotype assemblies. In addition, IGHV1-5, 3-7, 3-6, 4-3, and two IGHV6 genes were only identified in the primary fScoMax1.1 haplotype. All IGHV genes annotated in alternate fScoMax1.1 could be matched to a counterpart in the main genome assembly with more than 98% nucleotide sequence identity, suggesting that they can represent allelic variants. However, the analysis of ASM2237912v1 resulted in the identification of 2 IGHV genes, belonging to subgroups 3 and 1, which shared 96,3% nucleotide sequence identity with their fScoMax1.1 counterparts, indicating they can represent new genes (**Figure 1 and Supplementary File 1**).

The alignment of turbot IGHV amino acid sequences according to the IMGT unique numbering for V domain (Lefranc et al., 2005) (**Figure 5**) showed that all functional IGHV genes have identical FR lengths, regardless of the subgroup, while they present different CDR lengths. All members of IGHV1 and IGHV4 subgroups present [8.8.2] and [8.8.3] CDR lengths, respectively. IGHV2 and IGHV5, which are unique member subgroups, present [7.8.2] and [6.7.3] CDR lengths, respectively. The three IGHV genes belonging to subgroup 6 presented 100% sequence identity with [8.9.3] CDR lengths **(Figure 3 and 5)**. The largest subgroup, the IGHV3, has members with different CDR lengths, being [7.6.2] the most frequent, found in 7 out of 9 IGHV3 genes. The CDR lengths [7.7.2] and [6.6.2] are found in IGHV3-5 and in the IGHV3-2 genes, respectively. In addition, variants of IGHV3-1 and IGHV3-2 genes with different CDR lengths were identified in the ASM2237912v1 assembly.

**Figure 5.**
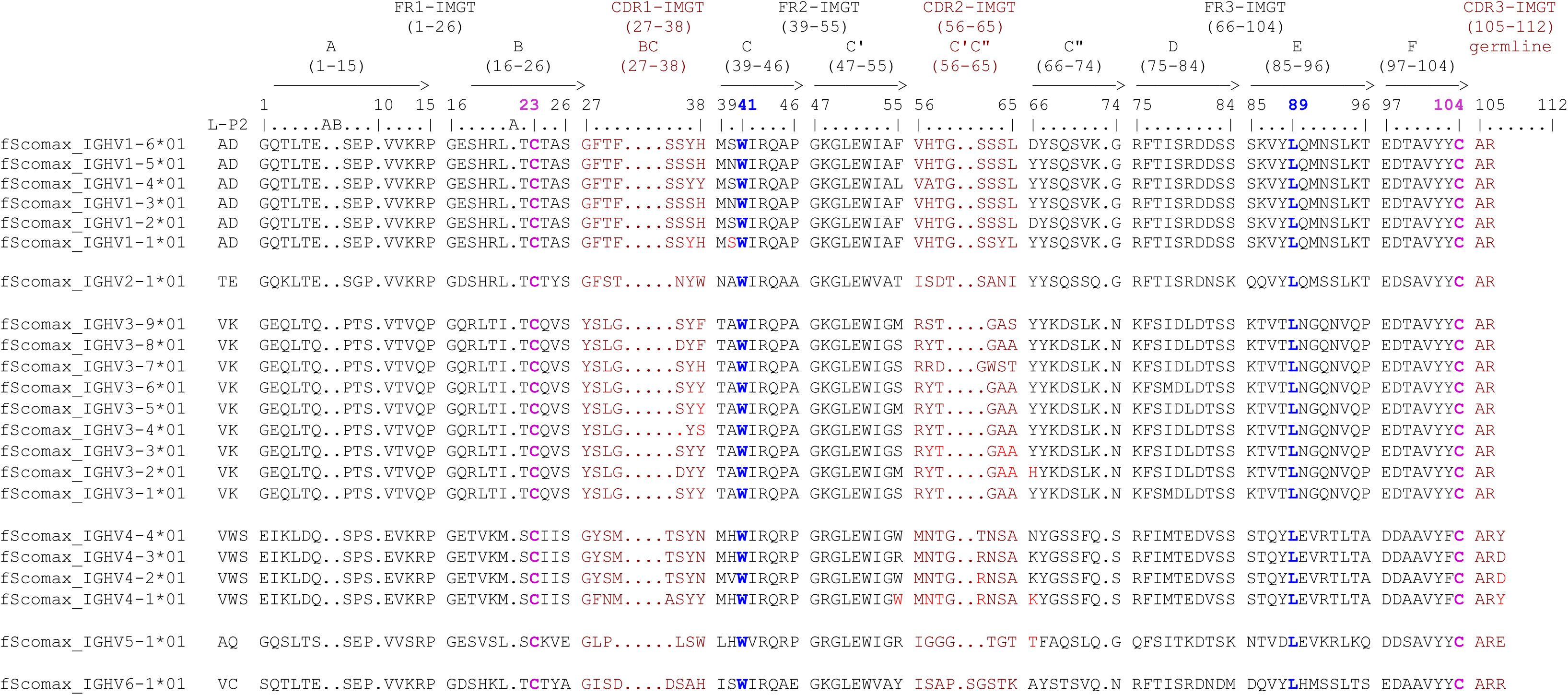
IMGT Protein display for turbot IGHV functional genes. The description of the strands and loops is according to the IMGT unique numbering for V-REGION (Lefranc et al., 2005, 2003). 1st-CYS C23, CONSERVED-TRP W41, hydrophobic L89 and 2nd-CYS C104 are colored (IMGT color menu) and in bold. Letters in red indicate aminoacid changes observed in IGHV genes annotated in the ASM2237912v1 and/or alternate haplotype of fScoMax1.1 genome assemblies.

### Expressed IGH Repertoire

The expression of turbot IGH genes was further characterized by deep sequencing of 5’-RACE products obtained from spleen and skin of four unvaccinated juvenile turbot. The 5’-RACE was initiated using IgM, IgD or IgT -specific constant region primers (**Supplementary File 2**). The summary statistics from the MiSeq, Cutadapt, pRESTO and IgBLAST analyses are provided in **Supplementary File 5**. Functional V(D)J rearrangements, defined as in-frame sequences lacking stop codons and containing a complete V(D)J junction, were considered for further analysis **(Supplementary File 5).** The percentage of functional sequences differed across isotypes but was consistent between tissues. Across all specimens and tissues, IgM was the dominant isotype in terms of productive and complete V(D)J, accounting for the majority of the functional repertoire in both skin (80.2 ± 6.4%) and spleen (74.0 ± 3.8%). In contrast, the relative abundance of functional IgD and IgT encoding rearrangements differed between tissues, in spleen approximately the 18,3% of unique sequences corresponded to IgD, and 7,7% to IgT, whereas in skin, IgT represented 14.1 ± 7.0% of functional transcripts and IgD only 5.8 ± 1.6%. This observation suggests a preferential compartmentalization of IgT within mucosal and skin-associated tissues, alongside a higher representation of IgD in the systemic immune compartment.

The analysis of IGHV gene usage was performed among all productive sequences using IgBLAST (Ye et al., 2013)(Ye et al., 2013). The IGHV genes annotated in the primary haplotype of the fScoMax1.1 genome assembly were used as the reference dataset. IgBLAST enabled unambiguous assignment of the IGHV gene in 86 % of the submitted sequences, including the exception of the IGHV1-3/IGHV1-5 gene pair and IGHV6-1/6-2/6-3 genes, which could not be distinguished because they share identical nucleotide sequences, as shown in the identity matrix **(see Figure 3).** For approximately 14 % of the sequences, IgBLAST reported different assignments to closely related germline IGHV genes belonging to the same subgroup, due to the high nucleotide similarity among genes within these families. These sequences were therefore retained for subgroup-level analyses but not for the IGHV-IGHJ gene pairing analysis.

As shown in **Figure 6**, IGHV3 was the dominant subgroup in most repertoires, especially for IgM and IgD, where it consistently represented the main contributor in both skin and spleen. On average, IGHV3 accounted for more than 70% of sequences in both tissues, followed by IGHV1 subgroup, which represented approximately 15% and 22% of sequences, in skin and spleen, respectively. IgD showed a similar overall pattern, although skin IgD was more variable. In contrast, IgT showed a different profile. In spleen, IgT repertoires expressed IGHV3 and IGHV4 as the main subgroups, representing 46% and 29% of sequences, respectively. Furthermore, in skin, IGHV4 represented a major fraction of the repertoire, accounting for 45% of sequences overall and becoming dominant in several individuals, in contrast to its minimal contribution to IgM and IgD repertoire (**Figure 6)**. Minor IGHV gene subgroups (IGHV2, IGHV5, IGHV6) collectively contributed less than 15% across all samples. Overall, these data reveal isotype associated differences in IGHV subgroup usage, with IgT showing a particular IGHV4 usage, especially in skin.

**Figure 6.**
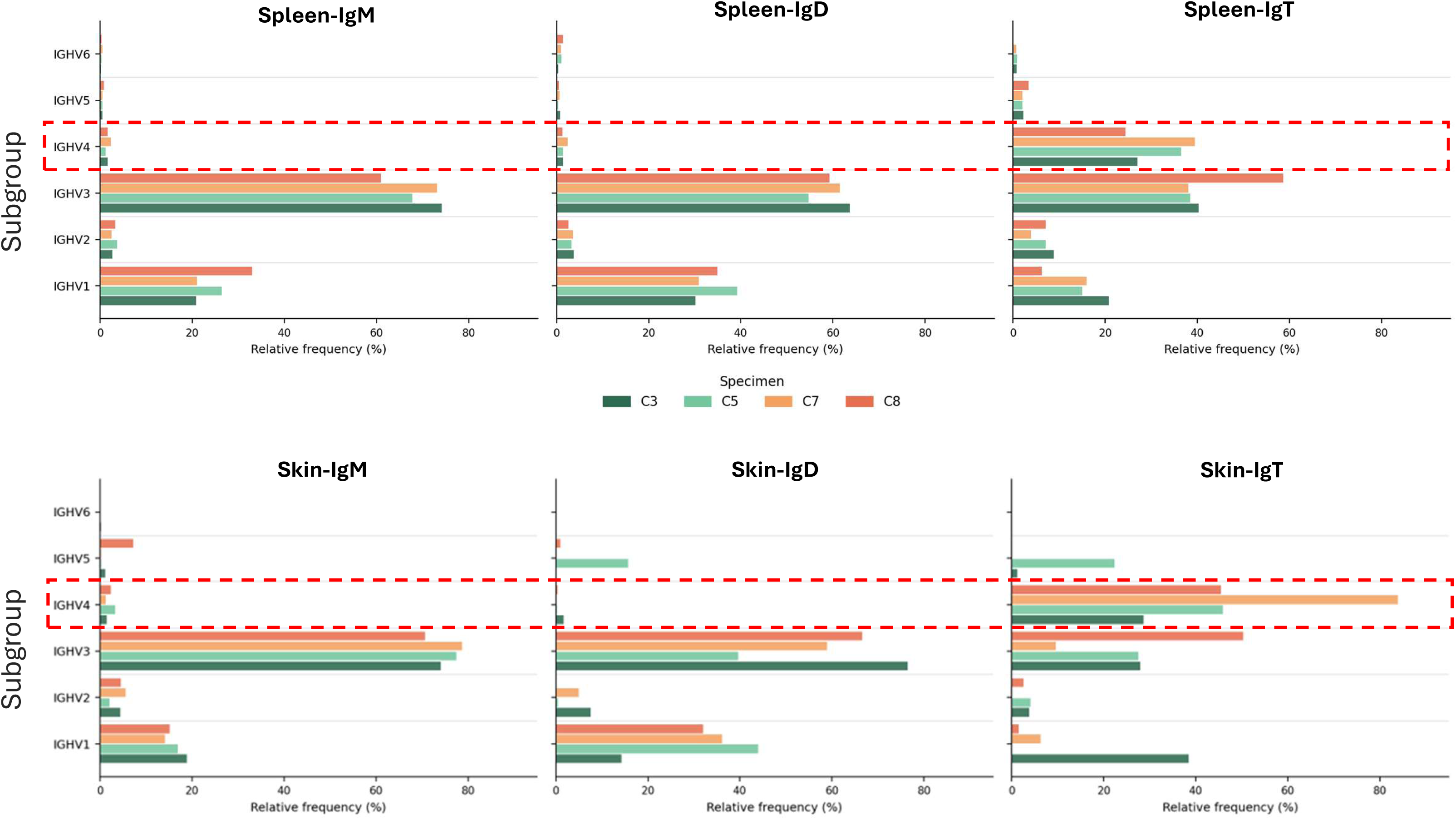
IGHV subgroup usage across IgM, IgD, and IgT repertoires in spleen and skin. Relative frequency of IGHV subgroup usage in productive IgM, IgD, and IgT sequences from spleen and skin of individual turbot specimens. Bars represent the percentage of sequences assigned to each IGHV subgroup within each specimen, tissue, and isotype. IgM and IgD repertoires were dominated by IGHV3 in both tissues, with a secondary contribution of IGHV1, whereas IGHV4 usage was limited in these isotypes. In contrast, IgT repertoires displayed a distinct IGHV profile, characterized by a marked contribution of IGHV4, particularly in skin. The dashed red boxes highlight the preferential usage of IGHV4 in IgT repertoires compared with IgM and IgD.

IGHV–IGHJ gene pairing analysis further supported these results (**Figure 7)**. The majority of annotated IGHV genes were identified within the IgM repertoire, which displayed a preferential usage of the IGHJ3 gene in both spleen and skin. IgD showed a similar IGHV-IGHJ pairing profile, while IgT displayed a different pairing pattern, with all sequences using IGHJT as the J gene segment. Being the strongest IgT signals observed for IGHV4-1 and IGHV3-8 in both skin and spleen. Several annotated IGHV genes showed little or no detectable contribution to the expressed repertoire. The low detection of IGHV1-2 and IGHV3-6 may reflect their high sequence identity with other members of the same subgroup (**Figure 3**), which could limit confident gene assignment. In contrast, the limited contribution of IGHV6-1 is more likely related to its association with a non-canonical recombination signal sequence, potentially reducing its rearrangement efficiency. As expected, IGHV(I), which was classified as a pseudogene due to the absence of the conserved Cys23 residue, was not detected in any tissue or isotype.

**Figure 7.**
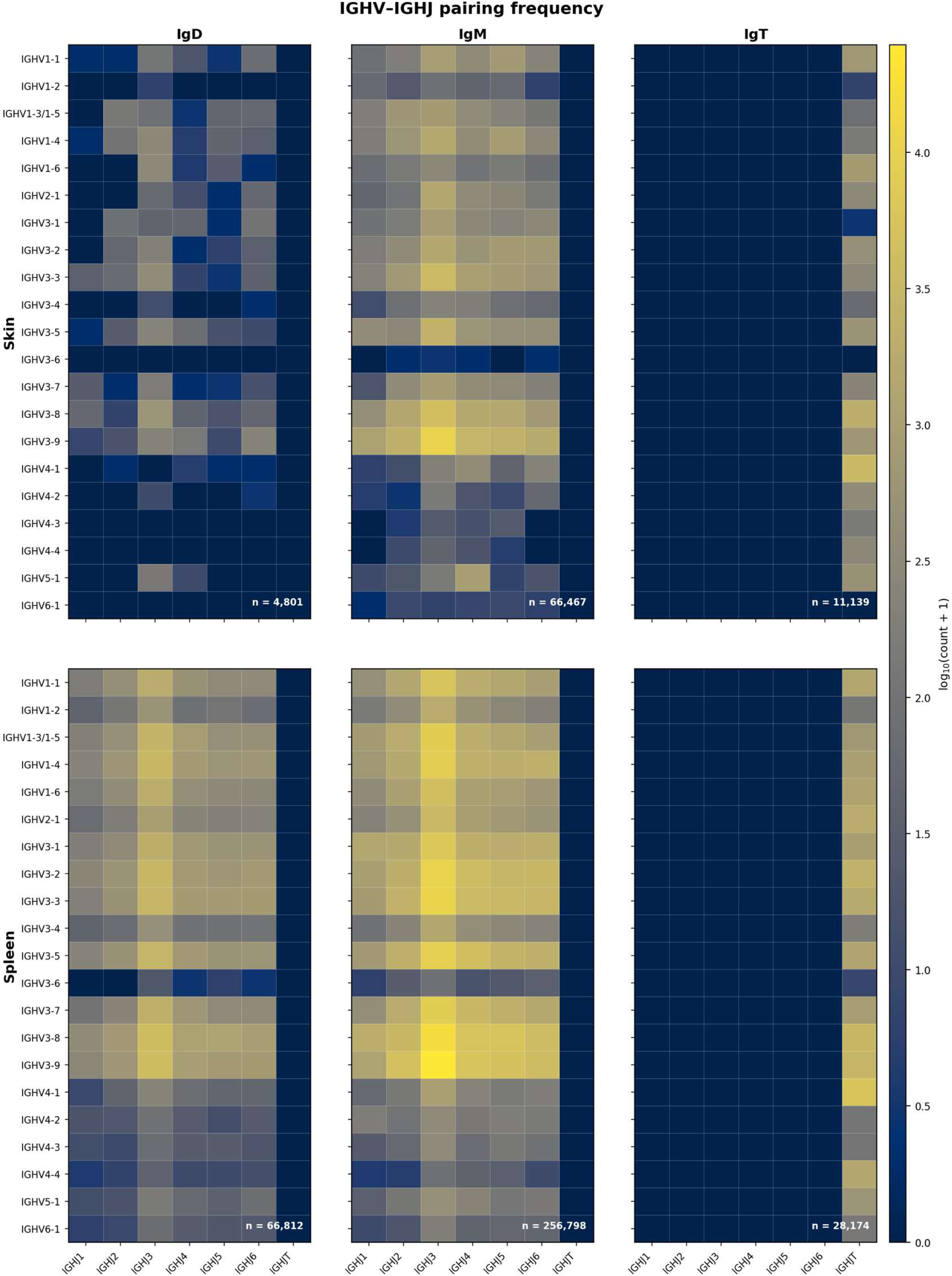
Turbot IGHV-IGHJ gene usage across tissues and immunoglobulin isotypes. Heatmaps showing combined IGHV-IGHJ pairing frequencies from four rainbow trout (C3, C5, C7, C8) in skin (top) and spleen (bottom) for IgD, IgM, and IgT isotypes. Color intensity represents log₁₀ (count + 1). IGHV genes are ordered by subgroup (IGHV1–IGHV6) and numerically within each family. Total productive unambiguous sequence counts per panel are indicated (n).

### Clonotype diversity

To further characterize the turbot antibody repertoire, productive sequences were analysed at the clonotype level. Clonotypes were defined by the combination of IGHV gene, IGHJ gene, and CDR3 nucleotide sequence, and repertoire coverage was assessed by rarefaction analysis. Rarefaction curves, generated by plotting the number of unique clonotypes against the number of subsampled consensus sequences, revealed marked differences in clonotype richness between tissues and isotypes (**Supplementary File 6**). Skin repertoires reached saturation rapidly, indicating that most clonotypes present in the sampled skin compartment were captured. In contrast, splenic repertoires showed greater clonotype richness and did not reach saturation, particularly for IgM, consistent with the larger and more diverse splenic B cell compartment. Rank abundance analysis supported these diversity patterns and further revealed pronounced tissue and isotype dependent differences in clonal dominance (**Figure 8**). Rank-abundance analysis showed that IgM was the least clonally dominated isotype in both tissues, consistent with a broad and highly diverse repertoire. The most abundant IgM clonotype represented only 1.3–5.0% of skin sequences and 0.3–0.6% of spleen sequences. Skin IgD repertoires were markedly more restricted than splenic IgD repertoires, with dominant clonotypes reaching up to 14.0%. IgT exhibited a clonal dominance in both tissues, but particularly in skin, where the leading clonotype represented 14–19% of sequences, consistent with a highly focused mucosal IgT response.

**Figure 8.**
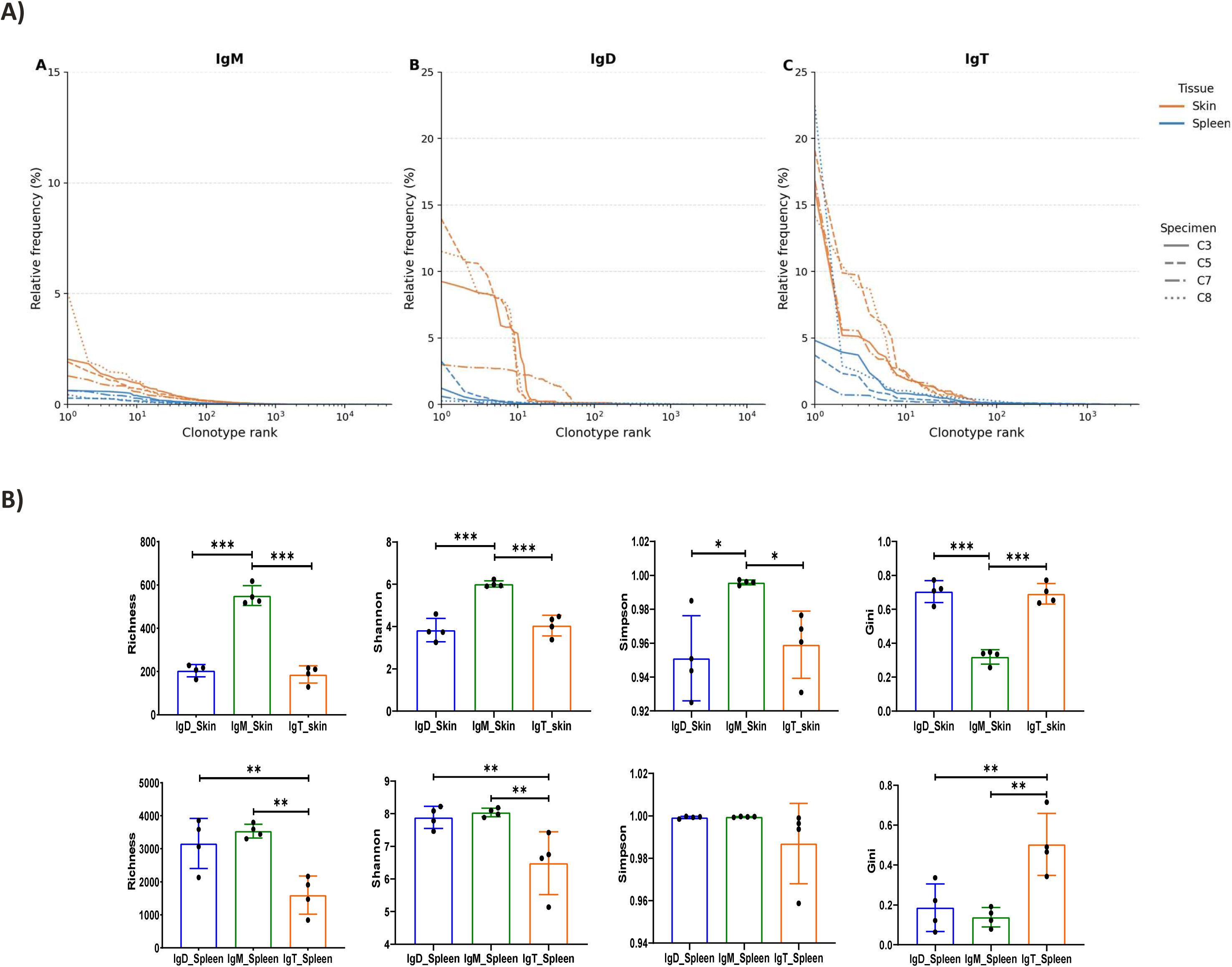
Turbot IG clonotype diversity. **A)** Rank abundance -Whittaker plots of clonotype repertoires in turbot skin and spleen. Clonotypes were defined by the combination of IGHV gene, IGHJ gene, and exact CDR3 nucleotide sequence, and restricted to productive sequences with complete V(D)J alignments. Each line represents one individual (n = 4 specimens; C3, C5, C7, C8; distinguished by line style: solid, dashed, dash-dot, and dotted, respectively). Orange lines: skin; blue lines: spleen. The x-axis shows clonotype rank in descending order of abundance (log scale); the y-axis shows the relative frequency of each clonotype within its library (linear scale). IgM y-axis is capped at 15%; IgD and IgT at 25%. . **B)** Comparison of clonotype diversity index among immunoglobulin isotypes and tissues. Diversity metrics were calculated after rarefaction to normalize for sequencing depth differences (Skin: 843 sequences; Spleen: 3583 sequences). Values are presented as mean ± standard deviation (n=4 fish per group). Richness represents the number of unique clonotypes observed. Shannon index measures diversity accounting for both richness and evenness. Simpson index represents the probability that two randomly selected sequences belong to different clonotypes. Gini coefficient measures clonal inequality (0 = perfectly even distribution; 1 = single dominant clone). P-values from one-way ANOVA comparing IgD, IgM, and IgT within each tissue, followed by Tukey’s Honest Significant Difference (HSD) post-hoc test for pairwise comparisons when ANOVA was significant (p<0.05). Significance levels: *p<0.05, **p<0.01, ***p<0.001.

To further compare immunoglobulin repertoire diversity, we analysed complementary diversity and clonality metrics, including Shannon, Simpson, and Gini indices (**Figure 8**). To minimize biases associated with differences in sequencing depth, datasets were normalized by subsampling based on the rarefaction curves and the smallest individual consensus sequence count within each tissue: 879 sequences for skin and 4,135 sequences for spleen. In skin, IgM showed significantly greater diversity than both IgD and IgT across the analysed metrics, including clonotype richness (one-way ANOVA, p < 0.001), whereas IgD and IgT did not differ significantly from each other (p = 0.80). In spleen, IgM and IgD displayed comparable diversity levels, while IgT showed a marked reduction relative to both isotypes, indicating a more restricted clonal composition of the systemic IgT repertoire.

### Tissue compartmentalization

In vertebrates, the tissue distribution and trafficking of B-cell clones are thought to play an important role in local immune protection and immune homeostasis (Lindner et al., 2015, 2012; Meng et al., 2017; Rhee et al., 2004), although the extent to which B-cell clones are shared between systemic and mucosal compartments remains poorly understood. To assess the degree of B cell clonotype overlap between skin and spleen in *Scophthalmus maximus*, we quantified clonotype sharing between both tissues using Jaccard and Morisita-Horn indices, which were calculated across the full IgM, IgD or IgT clonotype repertoires, and complemented by analysis of the 25 most abundant clonotypes (Top25) per library.

At the full-repertoire level, Jaccard indices were consistently low across all specimen and isotype combinations (**Figure 9**), indicating that most clonotype identities were tissue-restricted. This pattern was particularly evident in specimen C5, in which no overlap was detected for any of the three isotypes. Mean Jaccard values were similarly low for IgM, IgD, and IgT, reaching 0.016, 0.001, and 0.017, respectively. These low values were likely influenced by the different size between compartments, as spleen libraries were 7- to 75-fold larger than their matched skin libraries (Supplementary File 4). In contrast, Morisita–Horn indices, which account for clonotype abundance, revealed higher repertoire similarity for IgT (mean: 0.216) and IgM (mean: 0.110), whereas IgD overlap remained close to zero (**Figure 9**). Thus, although most clonotype identities were compartmentalized between skin and spleen, expanded IgM and, especially, IgT clones contributed to tissue sharing, suggesting trafficking of expanded B-cell clones.

**Figure 9.**
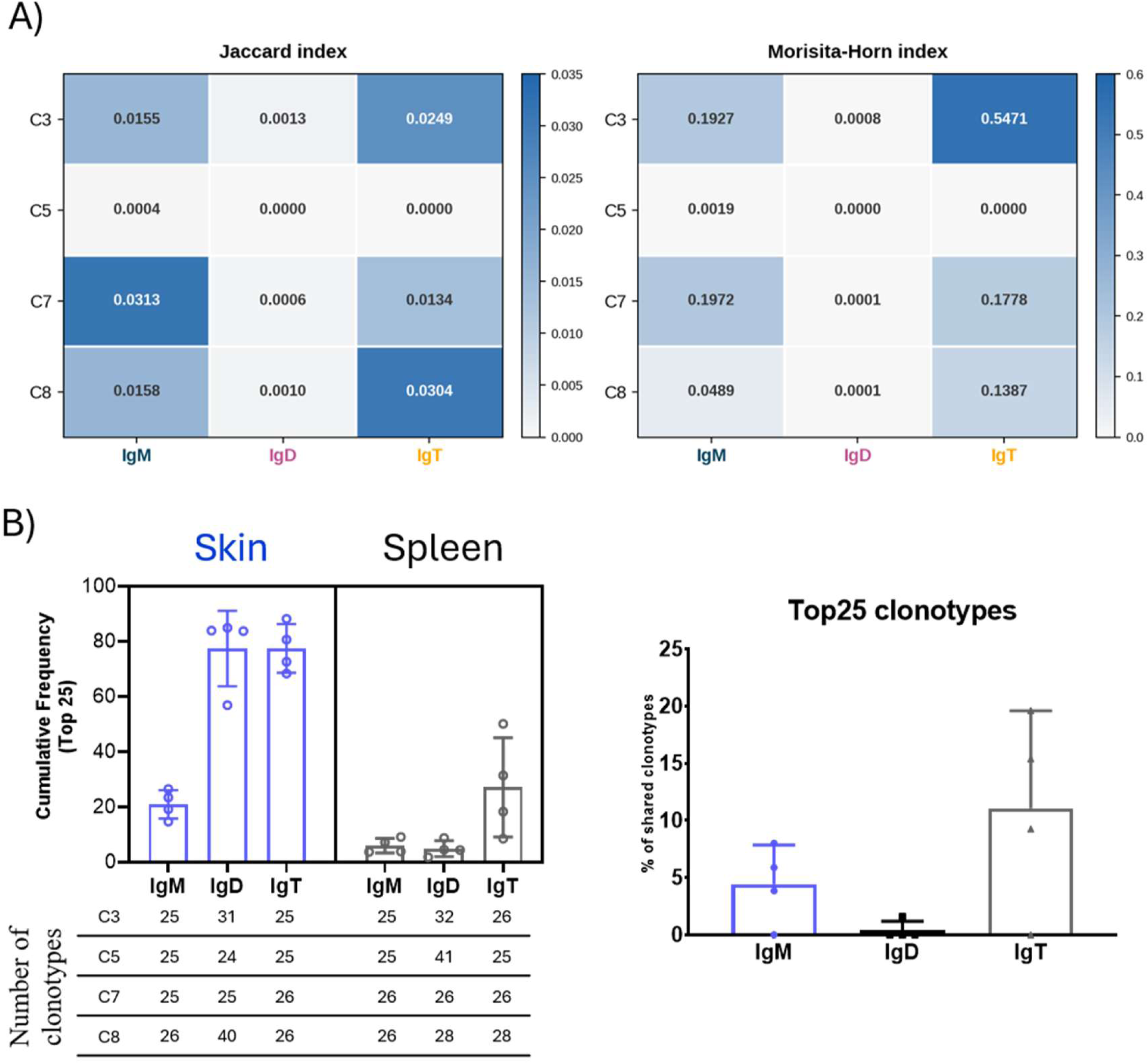
Clonotype sharing between spleen and skin across IgM, IgD, and IgT repertoires in turbot. **(A)** Heatmaps showing clonotype overlap between matched skin and spleen samples for each specimen and isotype, calculated across the full productive clonotype repertoire using the Jaccard and Morisita–Horn indices. Jaccard values quantify the fraction of shared clonotype identities between tissues, whereas Morisita–Horn values account for clonotype abundance. **(B)** Analysis of the 25 most abundant clonotypes per library. Left panel shows the cumulative frequency of the Top25 clonotypes in skin and spleen repertoires, with the number of clonotypes included for each specimen indicated below, reflecting ties at rank 25. Right panel shows the percentage of shared clonotypes among the combined Top25 skin and spleen clonotype sets for each isotype.

Analysis of the Top25 clonotypes further supported this isotype dependent pattern of tissue sharing. The cumulative frequency of these clonotypes, used as a measure of clonal dominance (Figure 9). Despite strong clonal dominance in both skin IgT and IgD repertoires, only dominant IgT clonotypes showed appreciable overlap with spleen, whereas dominant IgD clonotypes were largely tissue-restricted (Figure 9). Skin IgM displayed an intermediate pattern, with lower clonal dominance and moderate sharing. Together, these results indicate that clonal expansion alone does not predict tissue sharing. Rather, the relationship between dominance and tissue overlap differs among isotypes, with IgT showing the strongest evidence of shared dominant clonotypes between mucosal and systemic compartments.

## Discussion

In this study, we provide a comprehensive annotation of the turbot IGH locus and the use of this genomic framework to characterize the expressed antibody heavy-chain repertoire across systemic and skin-associated immune compartments. Analysis of the primary haplotype fScoMax1.1 assembly identified a compact IGH locus of approximately 72 kb on chromosome 19, containing 25 functional IGHV genes, one IGHV pseudogene, three IGHD genes, seven IGHJ genes, and three constant region genes corresponding to IgT, IgM and IgD. This organization is consistent with the translocon configuration described in other teleost species (Bao et al., 2010; Bradshaw and Valenzano, 2020; Danilova et al., 2005; Gambón-Deza et al., 2010; Hansen et al., 2005), while also revealing features specific to turbot, including a limited number of D segments and only one IGHJT gene. The conservation of the IGHD, IGHJ and IGHC gene content across the three analyzed assemblies indicates that the core D–J–C region of the turbot IGH locus is structurally stable, whereas variation in IGHV gene number and position suggests that the IGHV region represents the most dynamic component of this locus.

Differences in IGHV gene content among the primary and alternate fScoMax1.1 haplotypes and the ASM2237912v1 assembly may reflect both technical and biological factors. Adaptive antigen receptor loci are difficult to assemble because of their repetitive and highly similar sequences, and assembly fragmentation or haplotype collapse can affect accurate reconstruction of IGHV gene clusters. However, the CloseRead analysis supported high-confidence assembly of most annotated genes, particularly in the haplotype-resolved fScoMax1.1 assembly. Therefore, at least part of the observed IGHV variation may correspond to structural variation among turbot genomes. Such variation has been increasingly recognized as a relevant feature of antigen receptor loci in vertebrates, including mice and humans, where they influence the available germline repertoire (Rodriguez et al., 2023; Watson et al., 2019). In turbot, this may have consequences for inter-individual differences in antibody responses, although additional haplotype-resolved assemblies will be required to define the full extent of IGH locus polymorphism in this species.

The functional turbot IGHV genes annotated in the three analysed genome assemblies grouped into six subgroups, with a marked expansion of IGHV3 and, to a lesser extent, IGHV1 and IGHV4. This subgroup organization, supported by nucleotide identity (>75%) and phylogenetic analyses, provides a coherent reference dataset for repertoire annotation. Nevertheless, the presence of highly similar or nearly indistinguishable IGHV genes highlights an important limitation for gene-level assignment in expressed repertoire studies. Ambiguous assignment among closely related genes required conservative interpretation at the subgroup level, emphasizing the importance of combining genomic annotation with expressed repertoire data, particularly in species for which germline antigen receptor databases are newly established (Magadan et al., 2019b).

The analysis of productive V(D)J transcripts revealed clear isotype- and tissue-associated differences in the turbot IGH repertoire. IgM represented the dominant productive isotype in both skin and spleen, consistent with its important role at mucosal and systemic level in teleosts (Bilal et al., 2021; Ding et al., 2025; Kong et al., 2025). However, IgT represented a larger fraction of the productive repertoire in skin than in spleen, whereas IgD was more represented in spleen. This pattern agrees with the proposed specialization of IgT in mucosal tissues (Xu et al., 2016, 2013; Zhang et al., 2021, 2010). At the same time, the presence of substantial IgM expression in skin also indicates that mucosal antibody responses in turbot are unlikely to be mediated exclusively by IgT. Because the 5′-RACE approach using isotype-specific constant region primers measures expressed transcripts rather than absolute B cell frequencies; these differences should be interpreted as reflecting the expressed repertoire rather than the cellular composition of each compatment. In addition, the relative contribution of IgM and IgT to mucosal immunity may depend on species, tissue, developmental stage, and immune status (Faílde et al., 2014; Parra et al., 2015; Sheng et al., 2019).

The analysis of IGHV subgroup usage further supported an isotype specific organization of expressed repertoire. IgM and IgD showed broadly similar profiles, dominated by IGHV3 followed by IGHV1 in skin and spleen. This similarity is consistent with the developmental relationship between IgM- and IgD-expressing B cells, and with the structure of teleost IgD transcripts, which are generated by splicing between the Cμ1 exon and the Cδ1 exon, which results in a chimeric molecule (Fillatreau et al., 2013). In contrast, IgT displayed a distinct IGHV profile, particularly in skin, where IGHV4 accounted for a major fraction of the repertoire and became dominant in several individuals. The IGHV–IGHJ pairing analysis reinforced this distinction: IgM and IgD displayed broad pairing patterns with a marked contribution of IGHJ3, whereas IgT sequences paired exclusively with IGHJT, as expected from the organization of the turbot IGH locus. The strongest IgT signals involved IGHV4-1 and IGHV3-8 in both tissues, spleen and skin, suggesting that the IgT repertoire is characterized not only by exclusive IGHJT use but also by preferential recruitment of particular IGHV genes (Castro et al., 2013). Together, these findings indicate that the IgT repertoire is shaped by constraints that differ from those acting on IgM and IgD, potentially involving locus architecture, recombination accessibility, lineage-specific selection, or antigen-driven expansion in the skin compartment.

At the clonotype level, the turbot IGH repertoire showed a clear hierarchy of diversity among isotypes and tissues. IgM exhibited the broadest and least clonally dominated repertoire, particularly in spleen, where rarefaction curves did not reach saturation and tens of thousands of clonotypes were detected. This pattern is consistent with a large and diverse systemic B cell compartment containing many low frequency clonotypes, as reported in other species (Györkei et al., 2024; Magadan et al., 2019a). IgD showed an intermediate profile, with relatively diverse repertoires in spleen but more restricted repertoires in skin. IgT was the most clonally restricted isotype, especially in skin, where dominant clonotypes accounted for a substantial fraction of sequences. Whether these expansions reflect responses to environmental microbiota, previous antigen exposure, homeostatic selection, or intrinsic features of the IgT compartment remains to be determined.

Analysis of skin spleen clonotype sharing further indicated that systemic and skin associated antibody repertoire are compartmentalized. Shared clonotypes represented only a minor fraction of the splenic repertoire, consistent with the greater diversity of spleen, and overall Jaccard indices remained low across isotypes. This comparison should therefore be considered with caution, because skin sampling represented less than one quarter of the total skin surface, whereas the entire spleen was collected to isolate the RNA. In this context we likely underestimate the full extent of skin and spleen clonotype sharing. Despite this, abundance weighted Morisita–Horn indices and Top25 clonotype analyses revealed that a subset of expanded IgM and, especially, IgT clonotypes was shared between compartments. The restricted overlap between skin and spleen may reflect local maintenance or expansion of B cell clones in the skin, selective recruitment from systemic pools, or a combination of both. Although the number of specimens was limited, these data provide an initial baseline for understanding antibody repertoire compartmentalization in turbot. In future studies, longitudinal sampling after infection or vaccination will be needed to determine whether antigen specific turbot clonotypes are generated locally, recruited from spleen, or shared between compartments during an active immune response.

Overall, this study establishes a curated annotation of the turbot IGH locus and links germline organization to expressed antibody repertoire diversity across tissues and isotypes. The results show that the turbot IGH locus contains a conserved D–J–C region but a more variable IGHV gene cluster, and that this germline architecture is reflected in isotype specific patterns of IGHV gene usage and IGHJ gene pairing. Additional haplotype-resolved genomes may reveal further IGHV alleles or genes that refine sequence assignments. Functionally, IgM forms a broad and diverse repertoire, particularly in spleen, whereas IgT is characterized by exclusive IGHJT use, preferential recruitment of selected IGHV genes, enrichment in skin, and stronger clonal restrictions. These findings provide a foundation for future studies of antigen-specific antibody responses in turbot and will be valuable for evaluating mucosal and systemic responses to infection and vaccination in this aquaculture relevant species.

**Table 1:**
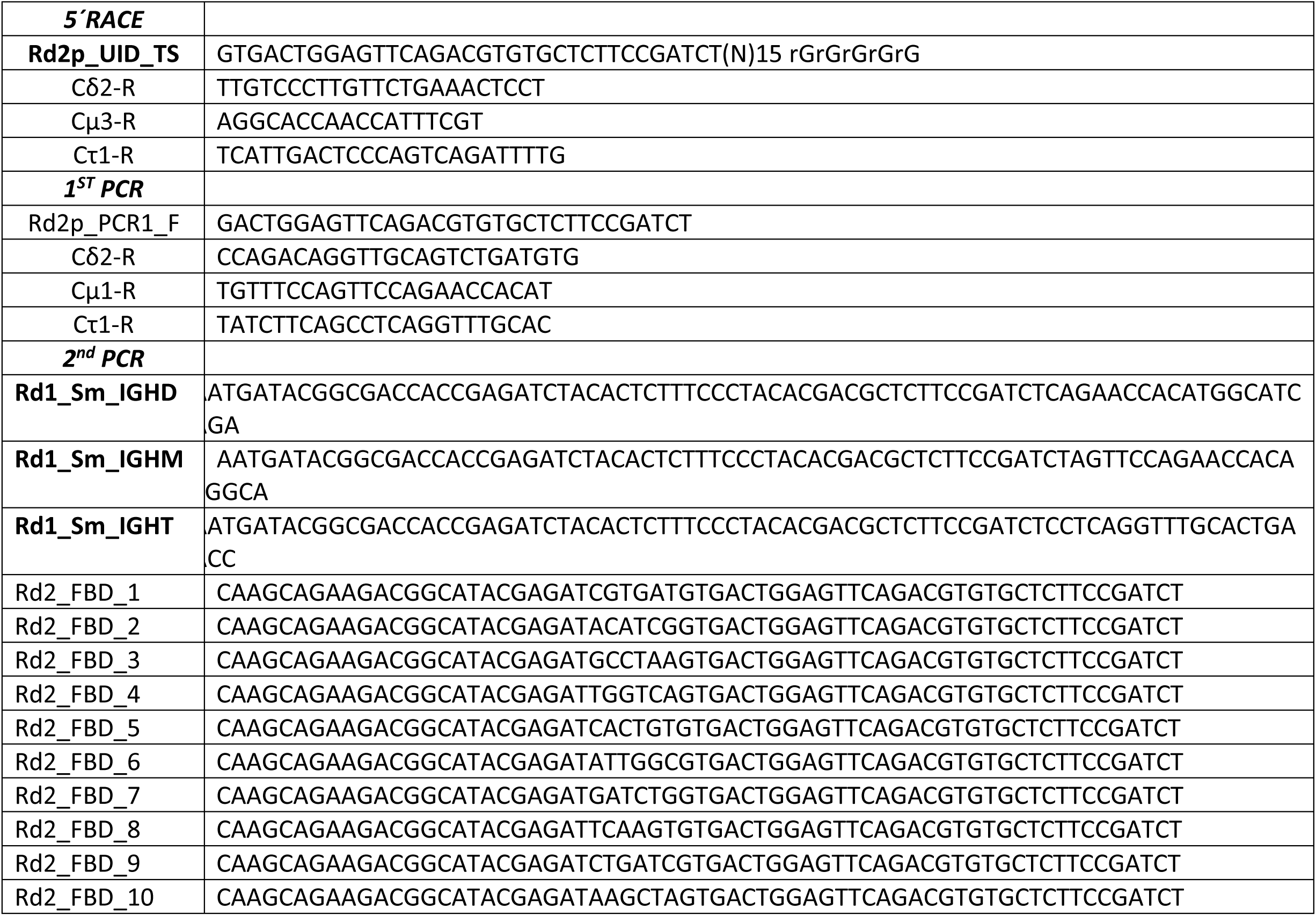
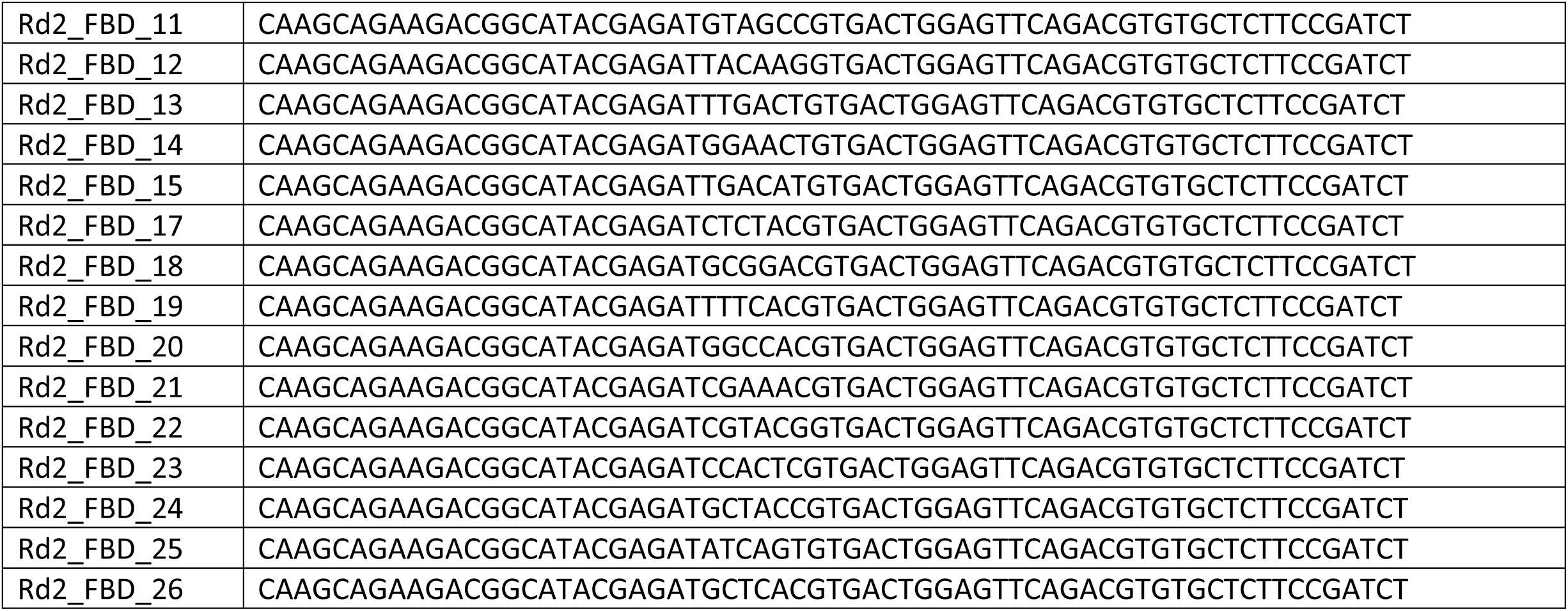
Oligos used to prepare amplicon libraries.

## Acknowledgment

We would like to thank Dr Corey Watson for his support in the data analysis performed using the CloseRead program, to thank Marie Paule Lefranc for her help with the locus annotation and Silvia Pérez for her support in the AIRR seq data analysis. This study was funded by the Agencia Estatal de Investigación (RODAVAC project; PID2022-137771OB-I00). RT was supported by a predoctoral fellowship from Xunta de Galicia (ED481A-2024-195). The founders had no role in study design, data collection and analysis, decision to publish, or preparation of the manuscript.

## Resource availability

Requests for further information and resources should be directed to and will be fulfilled by the lead contact, Susana Magadán.

## Author contributions

RT performed AIRR sequencing experiments and data analysis; YZ performed the CloseRead analysis; SMo performed annotation of IGH locus and experimental data; FG and PB contributed to data analysis and discussions. SM and YS conceptualization, data analysis, funding acquisition. SM wrote the original draft. All the authors reviewed and proofread the manuscript and the experimental results.

## Declaration of generative AI and AI-assisted technologies

During the preparation of this manuscript, the authors employed generative AI tools, including ChatGPT (OpenAI) and Gemini (Google), for language and code refinement. These tools were used solely to improve clarity, coherence, and style. All AI-assisted content was carefully reviewed and substantively edited by the authors, who accept full responsibility for the accuracy and integrity of the final submitted work.

## Supplementary Files

**Supplementary File 1:** Coordinates of annotated IGH genes across the three turbot genome assemblies analyzed. The figure summarizes the presence, genomic localization, and chromosomal distribution of IGH genes in each assembly (fScoMax1.1 primary haplotype and alternative haplotype, and the ASM2237912v1 assembly) and includes the percentage sequence identity among corresponding loci.

**Supplementary File 2:** Table of oligos used for the turbot IGH repertoire analysis.

**Supplementary File 3:** Turbot IGHD and IGHJ sequences. 3A) Comparison of turbot IGHD genes and RSS sequence. Sequence comparison of identified IGHD genes in the three turbot genome assemblies: fScoMax1.1 primary haplotype (fScomax), the fScoMax1.1 alternative haplotype (afScomax), and the ASM2237912v1 assembly (yScomax). 3B) Comparison of turbot IGHJ genes and RSS sequence. Sequence comparison of all annotated IGHJ genes in the three turbot genome assemblies: fScoMax1.1 primary haplotype (fScomax), the fScoMax1.1 alternative haplotype (afScomax), and the ASM2237912v1 assembly (yScomax). The identified splicing site is underlined.

**Supplementary File 4:** CloseRead results. Analysis of IGH locus quality in the primary turbot genome (ASM2237912v1) assembly. Summary statistics of the reads alignment showing mapping quality across IGH locus, Mismatch rates of reads, Number of reads with indels of consecutive length of at least 2 bp and count of soft clipped bases in each read are shown in different graphics. The IGH locus length is shown using blue bar charts. The detailed analysis of alignment mismatch includes read coverage across the entire IGH locus, color-coded by mapping quality, and base pair mismatch rate.

**Supplementary File 5:** Summary statistics of MiSeq and IgBlast results.

**Supplementary File 6:** Rarefaction analysis of IgM, IgD, and IgT clonotype richness in skin and spleen repertoires. Rarefaction curves showing the number of unique clonotypes detected as a function of the number of subsampled productive consensus sequences for each specimen. Clonotypes were defined by the combination of IGHV gene, IGHJ gene, and CDR3 nucleotide sequence. Curves are shown separately for skin and spleen IgD, IgM, and IgT repertoires. Skin repertoires, particularly IgD and IgT, approached saturation more rapidly than splenic repertoires, indicating lower clonotype richness and greater sampling coverage in the skin compartment. In contrast, spleen repertoires, especially IgM, showed markedly higher clonotype richness and did not reach saturation, consistent with a broader and more diverse systemic antibody repertoire.

---

**Supplementary Fig 3.**
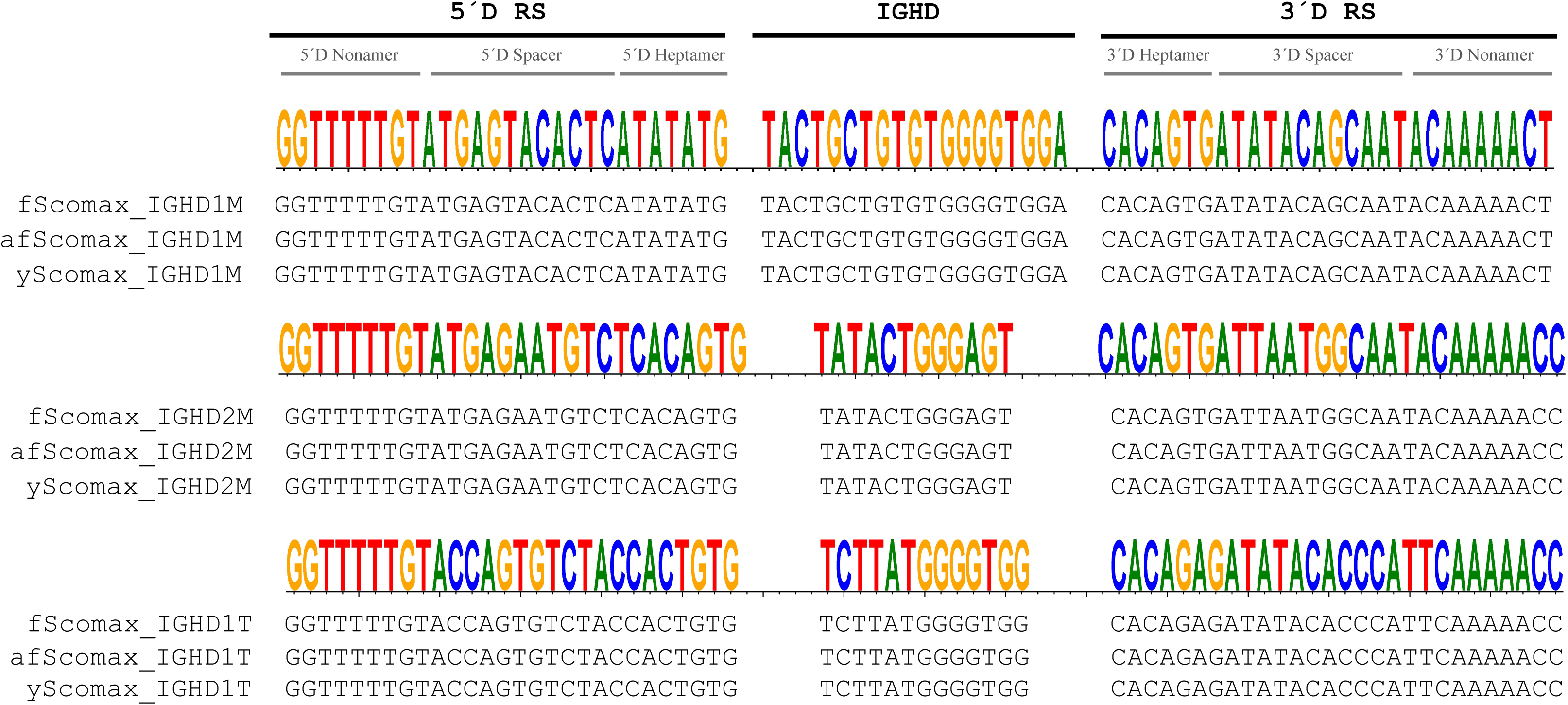
A: Comparison of turbot IGHD genes and RSS sequence. Sequence comparison of identified IGHD genes in the three turbot genome assemblies: fScoMax1.1 primary haplotype (fScomax), the fScoMax1.1 alternative haplotype (afScomax), and the ASM2237912v1 assembly (yScomax).

**Supplementary Fig 3.**
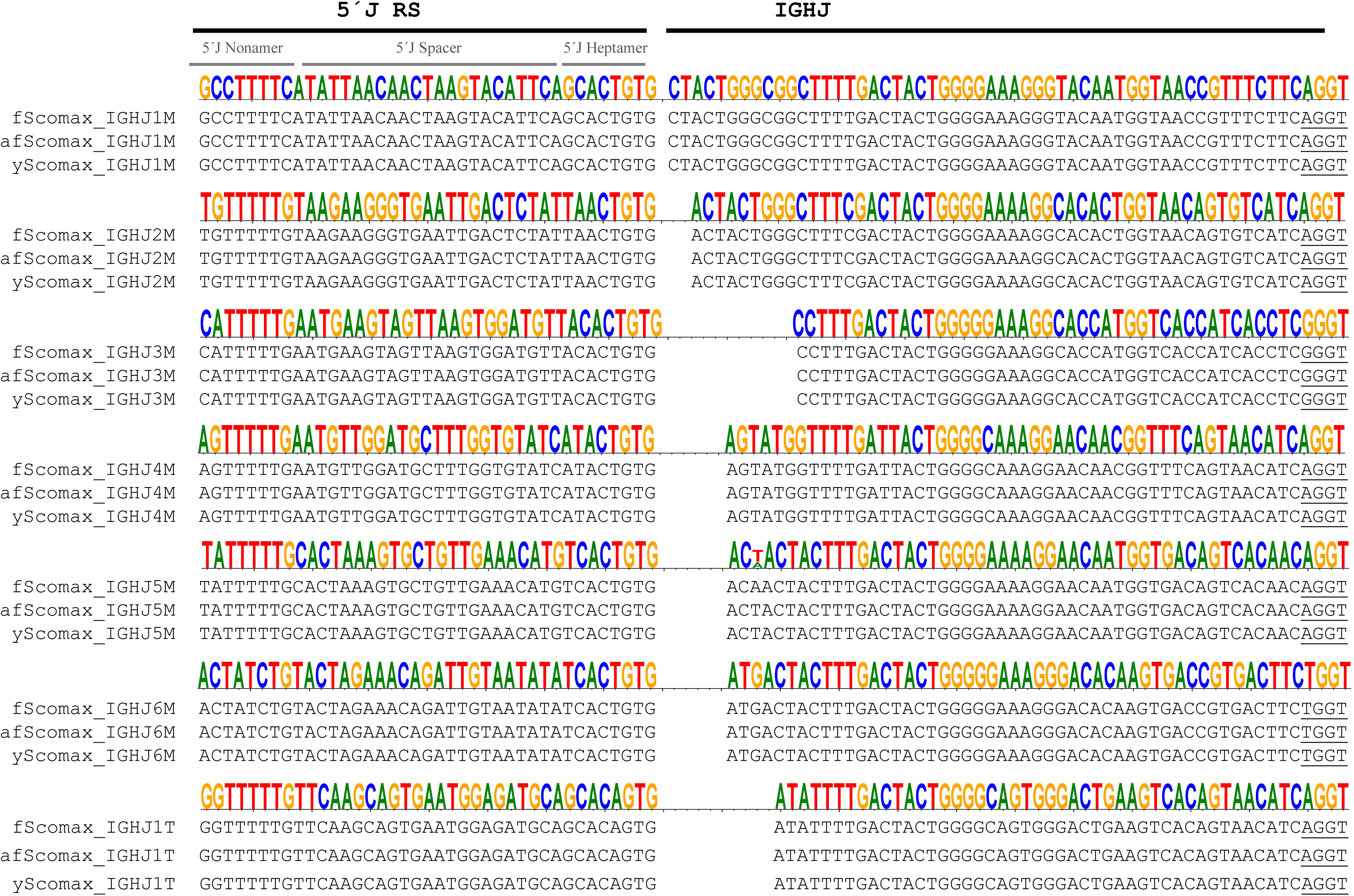
B: Comparison of turbot IGHJ genes and RSS sequence. Sequence comparison of all annotated IGHJ genes in the three turbot genome assemblies: fScoMax1.1 primary haplotype (fScomax), the fScoMax1.1 alternative haplotype (afScomax), and the ASM2237912v1 assembly (yScomax). The identified splicing site is underlined.

**Supplementary Figure 4:**
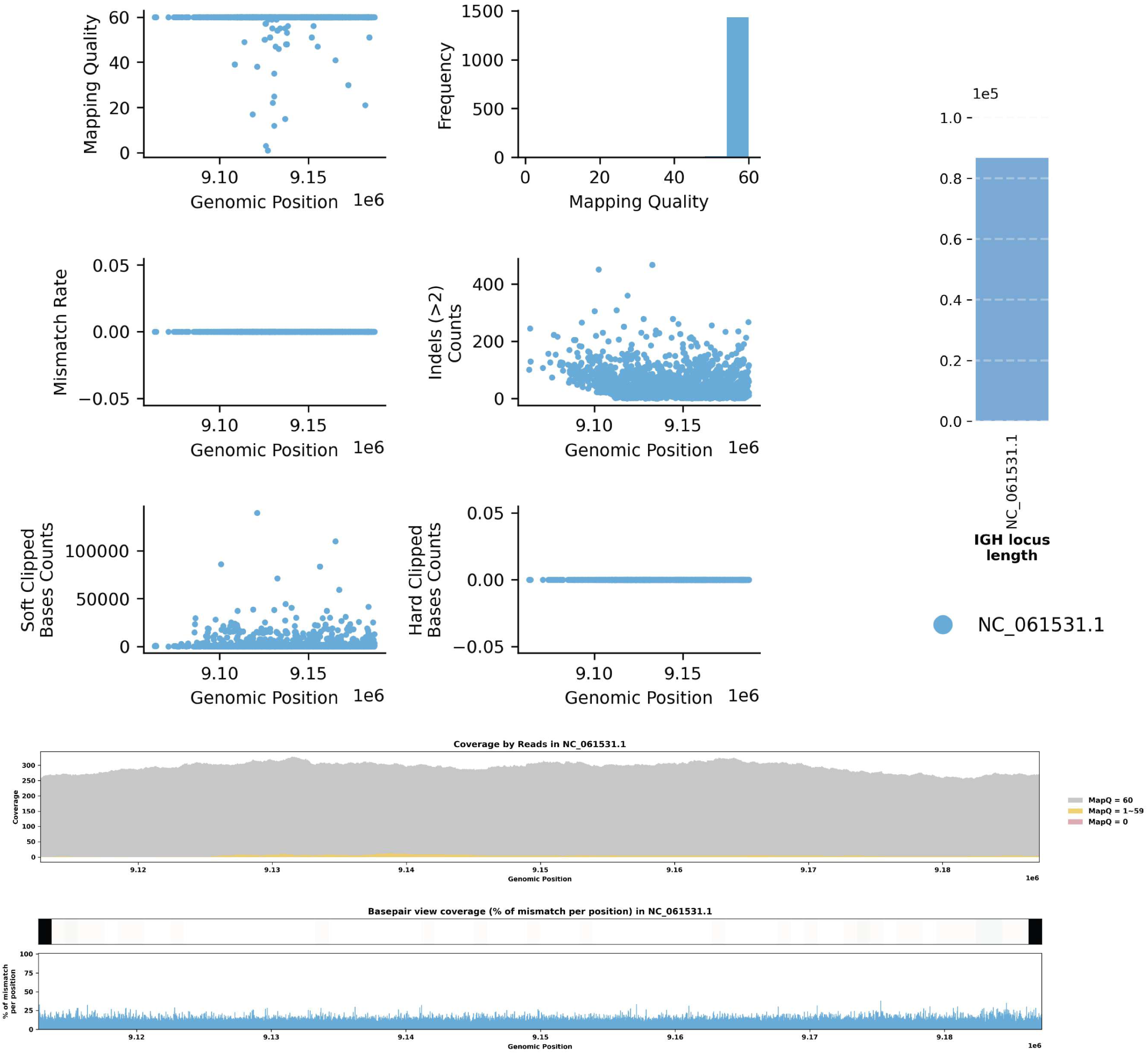
Detailed analysis of IGH locus assembly errors in the primary turbot genome assembly, ASM2237912v (GCA_022379125.1) Summary statistics of the read alignment, showing mapping quality across IGH loci. Mismatch rates of reads, Number of reads with indels of consecutive length of at least 2 bp, and count of soft clipped bases in each read, At the bottom, IGH locus length and detailed analysis of alignment mismatch in both haplotypes includes (i) read coverage across the entire IGH loci, color-coded by mapping quality, and (ii) basepair-oriented mismatch rate, a heatmap above indicating the frequency of high mismatch rate base pairs.

**Supplementary Fig 6.**
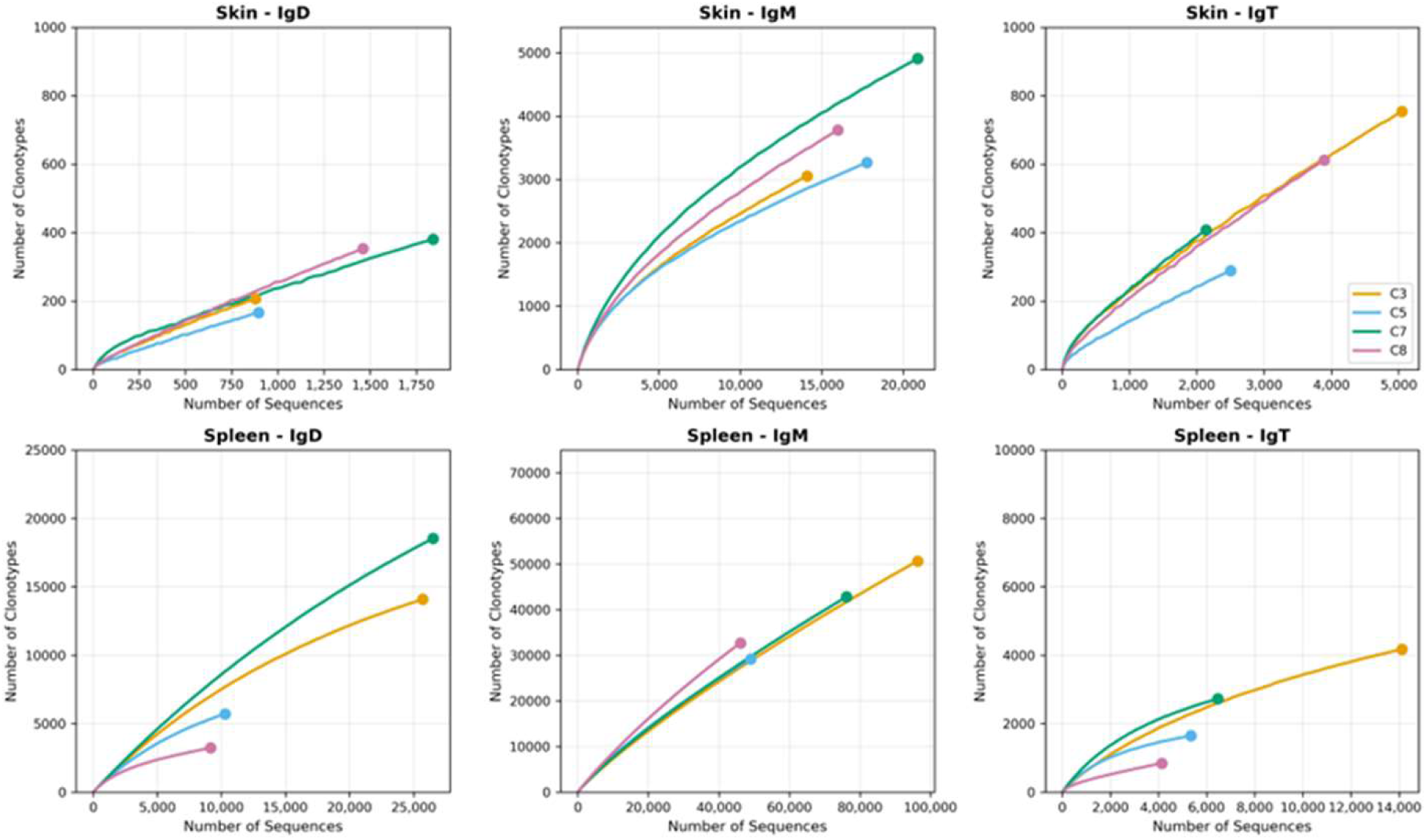
Rarefaction analysis of immunoglobulin repertoire diversity in turbot tissues. Rarefaction curves showing the relationship between sequencing depth (number of sequences) and clonotype richness (number of unique clonotypes) in Spleen and Skin tissues from four individual fish (C3, C5, C7, C8) distinguished by color.

## Bibliography

1. Al Hayek, S., Praité, A., Lambert, J.M., Bender, S., Delpy, L., 2025. 5’RACE Method for RNA-Based Sequencing of the Mouse Immunoglobulin Repertoire. Methods Mol. Biol. 2962, 103–119. 10.1007/978-1-0716-4726-4_8

2. Alt, F.W., Yancopoulos, G.D., Blackwell, T.K., Wood, C., Thomas, E., Boss, M., Coffman, R., Rosenberg, N., Tonegawa, S., Baltimore, D., 1984. Ordered rearrangement of immunoglobulin heavy chain variable region segments. EMBO J. 3, 1209–1219. 10.1002/J.1460-2075.1984.TB01955.X

3. Bao, Y., Wang, T., Guo, Y., Zhao, Z., Li, N., Zhao, Y., 2010. The immunoglobulin gene loci in the teleost Gasterosteus aculeatus. Fish Shellfish Immunol. 28, 40–48. 10.1016/j.fsi.2009.09.014

4. Bashford-Rogers, R.J.M., Bergamaschi, L., McKinney, E.F., Pombal, D.C., Mescia, F., Lee, J.C., Thomas, D.C., Flint, S.M., Kellam, P., Jayne, D.R.W., Lyons, P.A., Smith, K.G.C., 2019. Analysis of the B cell receptor repertoire in six immune-mediated diseases. Nature 574, 122–126. 10.1038/s41586-019-1595-3

5. Bengtén, E., Quiniou, S., Hikima, J., Waldbieser, G., Warr, G.W., Miller, N.W., Wilson, M., 2006. Structure of the catfish IGH locus: analysis of the region including the single functional IGHM gene. Immunogenetics 58, 831–844. 10.1007/s00251-006-0139-9

6. Bilal, S., Etayo, A., Hordvik, I., 2021. Immunoglobulins in teleosts. Immunogenetics 73, 65–77. 10.1007/s00251-020-01195-1

7. Boudinot, P., Novas, S., Jouneau, L., Mondot, S., Lefranc, M.P., Grimholt, U., Magadán, S., 2023. Evolution of T cell receptor beta loci in salmonids. Front. Immunol. 14. 10.3389/FIMMU.2023.1238321

8. Bradshaw, W.J., Valenzano, D.R., 2020. Extreme genomic volatility characterizes the evolution of the immunoglobulin heavy chain locus in cyprinodontiform fishes. Proceedings. Biol. Sci. 287. 10.1098/rspb.2020.0489

9. Brittain, R., Adkins, P., Scott-Somme, K., Modepali, V., 2025. The genome sequence of the turbot, Scophthalmus maximus (Linnaeus, 1758) (Pleuronectiformes: Scophthalmidae). Wellcome Open Res. 10, 592. 10.12688/wellcomeopenres.25070.1

10. Castro, R., Jouneau, L., Pham, H.-P., Bouchez, O., Giudicelli, V., Lefranc, M.-P., Quillet, E., Benmansour, A., Cazals, F., Six, A., Fillatreau, S., Sunyer, O., Boudinot, P., 2013. Teleost Fish Mount Complex Clonal IgM and IgT Responses in Spleen upon Systemic Viral Infection. PLoS Pathog. 9, e1003098. 10.1371/journal.ppat.1003098

11. Chen, W., Huang, J., Wang, W., Wang, Y., Chen, H., Wang, Q., Zhang, Y., Liu, Q., Yang, D., 2022. Multi-tissue scRNA-seq reveals immune cell landscape of turbot (Scophthalmus maximus). Fundam. Res. 2, 550–561. 10.1016/j.fmre.2021.12.015

12. Community, G., 2024. The Galaxy platform for accessible, reproducible, and collaborative data analyses: 2024 update. Nucleic Acids Res. 52, W83–W94. 10.1093/NAR/GKAE410

13. Danilova, N., Bussmann, J., Jekosch, K., Steiner, L.A., 2005. The immunoglobulin heavy-chain locus in zebrafish: identification and expression of a previously unknown isotype, immunoglobulin Z. Nat. Immunol. 6, 295–302. 10.1038/ni1166

14. Ding, Y., Fernández-Montero, A., Mani, A., Casadei, E., Miyazawa, R., Zhou, C., Chaumont, L., Posavi, M., Cole, S.D., Shibasaki, Y., Takizawa, F., Salinas, I., Sunyer, J.O., 2025. Secretory IgM regulates gut microbiota homeostasis and metabolism. Nat. Microbiol. 10, 1431–1446. 10.1038/S41564-025-02013-8

15. Edholm, E.S., Fenton, C.G., Mondot, S., Paulssen, R.H., Lefranc, M.P., Boudinot, P., Magadan, S., 2021. Profiling the T Cell Receptor Alpha/Delta Locus in Salmonids. Front. Immunol. 12. 10.3389/FIMMU.2021.753960

16. Eugster, A., Bostick, M.L., Gupta, N., Mariotti-Ferrandiz, E., Kraus, G., Meng, W., Soto, C., Trück, J., Stervbo, U., Luning Prak, E.T., 2022. AIRR Community Guide to Planning and Performing AIRR-Seq Experiments. Methods Mol. Biol. 2453, 261–278. 10.1007/978-1-0716-2115-8_15

17. Faílde, L.D., Losada, A.P., Bermúdez, R., Santos, Y., Quiroga, M.I., 2014. Evaluation of immune response in turbot (Psetta maxima L.) tenacibaculosis: Haematological and immunohistochemical studies. Microb. Pathog. 76, 1–9. 10.1016/j.micpath.2014.08.008

18. Figueras, A., Robledo, D., Corvelo, A., Hermida, M., Pereiro, P., Rubiolo, J.A., Gómez-Garrido, J., Carreté, L., Bello, X., Gut, M., Gut, I.G., Marcet-Houben, M., Forn-Cuní, G., Galán, B., García, J.L., Abal-Fabeiro, J.L., Pardo, B.G., Taboada, X., Fernández, C., Vlasova, A., Hermoso-Pulido, A., Guigó, R., Álvarez-Dios, J.A., Gómez-Tato, A., Viñas, A., Maside, X., Gabaldón, T., Novoa, B., Bouza, C., Alioto, T., Martínez, P., 2016. Whole genome sequencing of turbot (Scophthalmus maximus; Pleuronectiformes): a fish adapted to demersal life. DNA Res. 23, 181–192. 10.1093/dnares/dsw007

19. Fillatreau, S., Six, A., Magadan, S., Castro, R., Sunyer, J.O., Boudinot, P., 2013. The Astonishing Diversity of Ig Classes and B Cell Repertoires in Teleost Fish. Front. Immunol. 4. 10.3389/fimmu.2013.00028

20. Flajnik, M.F., 2005. The last flag unfurled? A new immunoglobulin isotype in fish expressed in early development. Nat. Immunol. 6, 229–230. 10.1038/ni0305-229

21. Fournier-Betz, V., Quentel, C., Lamour, F., Leven, A., 2000. Immunocytochemical detection of Ig-positive cells in blood, lymphoid organs and the gut associated lymphoid tissue of the turbot (Scophthalmus maximus). Fish Shellfish Immunol. 10, 187–202. 10.1006/fsim.1999.0235

22. Gambón-Deza, F., Sánchez-Espinel, C., Magadán-Mompó, S., 2010. Presence of an unique IgT on the IGH locus in three-spined stickleback fish (Gasterosteus aculeatus) and the very recent generation of a repertoire of VH genes. Dev. Comp. Immunol. 34, 114–122. 10.1016/j.dci.2009.08.011

23. Gao, C., Fu, Q., Su, B., Zhou, S., Liu, F., Song, L., Zhang, M., Ren, Y., Dong, X., Tan, F., Li, C., 2016. Transcriptomic profiling revealed the signatures of intestinal barrier alteration and pathogen entry in turbot (Scophthalmus maximus) following Vibrio anguillarum challenge. Dev. Comp. Immunol. 65, 159–168. 10.1016/j.dci.2016.07.007

24. Gao, Y., Yi, Y., Wu, H., Wang, Q., Qu, J., Zhang, Y., 2014. Molecular cloning and characterization of secretory and membrane-bound IgM of turbot. Fish Shellfish Immunol. 40, 354–361. 10.1016/j.fsi.2014.07.011

25. Garzón-Ospina, D., Buitrago, S.P., 2020. Igh locus structure and evolution in Platyrrhines: new insights from a genomic perspective. Immunogenetics 72, 165–179. 10.1007/s00251-019-01151-8

26. Gomez, D., Sunyer, J.O., Salinas, I., 2013. The mucosal immune system of fish: The evolution of tolerating commensals while fighting pathogens. Fish Shellfish Immunol. 35, 1729–1739. 10.1016/j.fsi.2013.09.032

27. Györkei, Á., Johansen, F.E., Qiao, S.W., 2024. Systematic characterization of immunoglobulin loci and deep sequencing of the expressed repertoire in the Atlantic cod (Gadus morhua). BMC Genomics 25. 10.1186/S12864-024-10571-0

28. Hansen, J.D., Landis, E.D., Phillips, R.B., 2005. Discovery of a unique Ig heavy-chain isotype (IgT) in rainbow trout: Implications for a distinctive B cell developmental pathway in teleost fish. Proc. Natl. Acad. Sci. 102, 6919–6924. 10.1073/pnas.0500027102

29. Hsu, E., 2009. V(D)J recombination: of mice and sharks. Adv. Exp. Med. Biol. 650, 166–79.

30. Kong, W., Wang, X., Ding, G., Yang, P., Shi, Y., Cai, C., Yang, X., Cheng, G., Takizawa, F., Xu, Z., 2025. Newly discovered and conserved role of IgM against viral infection in an early vertebrate. Elife 14. 10.7554/ELIFE.104465

31. Kumar, S., Stecher, G., Tamura, K., 2016. MEGA7: Molecular Evolutionary Genetics Analysis Version 7.0 for Bigger Datasets. Mol. Biol. Evol. 33, 1870–1874. 10.1093/MOLBEV/MSW054

32. Lefranc, M.-P., Lefranc, G., 2001. The immunoglobulin factsbook. Academic Press.

33. Lefranc, M.-P., Pommié, C., Kaas, Q., Duprat, E., Bosc, N., Guiraudou, D., Jean, C., Ruiz, M., Da Piédade, I., Rouard, M., Foulquier, E., Thouvenin, V., Lefranc, G., 2005. IMGT unique numbering for immunoglobulin and T cell receptor constant domains and Ig superfamily C-like domains. Dev. Comp. Immunol. 29, 185–203. 10.1016/j.dci.2004.07.003

34. Lefranc, M.P., 2014. Immunoglobulin and T cell receptor genes: IMGT® and the birth and rise of immunoinformatics. Front. Immunol. 5. 10.3389/fimmu.2014.00022

35. Lewis, S.M., 1994. The mechanism of V(D)J joining: Lessons from molecular, immunological, and comparative analyses. Adv. Immunol. 56, 27–150. 10.1016/s0065-2776(08)60450-2

36. Lindner, C., Thomsen, I., Wahl, B., Ugur, M., Sethi, M.K., Friedrichsen, M., Smoczek, A., Ott, S., Baumann, U., Suerbaum, S., Schreiber, S., Bleich, A., Gaboriau-Routhiau, V., Cerf-Bensussan, N., Hazanov, H., Mehr, R., Boysen, P., Rosenstiel, P., Pabst, O., 2015. Diversification of memory B cells drives the continuous adaptation of secretory antibodies to gut microbiota. Nat. Immunol. 16, 880–888. 10.1038/ni.3213

37. Lindner, C., Wahl, B., Föhse, L., Suerbaum, S., Macpherson, A.J., Prinz, I., Pabst, O., 2012. Age, microbiota, and T cells shape diverse individual IgA repertoires in the intestine. J. Exp. Med. 209, 365–377. 10.1084/jem.20111980

38. Magadán-Mompó, S., Sánchez-Espinel, C., Gambón-Deza, F., 2011. Immunoglobulin heavy chains in medaka (Oryzias latipes). BMC Evol. Biol. 11, 165. 10.1186/1471-2148-11-165

39. Magadan, S., Jouneau, L., Boudinot, P., Salinas, I., 2019a. Nasal Vaccination Drives Modifications of Nasal and Systemic Antibody Repertoires in Rainbow Trout. J. Immunol. ji1900157. 10.4049/jimmunol.1900157

40. Magadan, S., Krasnov, A., Hadi-Saljoqi, S., Afanasyev, S., Mondot, S., Lallias, D., Castro, R., Salinas, I., Sunyer, O., Hansen, J., Koop, B.F., Lefranc, M.P., Boudinot, P., 2019b. Standardized IMGT® Nomenclature of Salmonidae IGH Genes, the Paradigm of Atlantic Salmon and Rainbow Trout: From Genomics to Repertoires. Front. Immunol. 10. 10.3389/FIMMU.2019.02541

41. Magadan, S., Sunyer, O.J., Boudinot, P., 2015. Unique Features of Fish Immune Repertoires: Particularities of Adaptive Immunity Within the Largest Group of Vertebrates. pp. 235–264. 10.1007/978-3-319-20819-0_10

42. Meng, W., Zhang, B., Schwartz, G.W., Rosenfeld, A.M., Ren, D., Thome, J.J.C., Carpenter, D.J., Matsuoka, N., Lerner, H., Friedman, A.L., Granot, T., Farber, D.L., Shlomchik, M.J., Hershberg, U., Luning Prak, E.T., 2017. An atlas of B-cell clonal distribution in the human body. Nat. Biotechnol. 35, 879–886. 10.1038/NBT.3942

43. Parra, D., Korytář, T., Takizawa, F., Sunyer, J.O., 2016. B cells and their role in the teleost gut. Dev. Comp. Immunol. 64, 150–166. 10.1016/j.dci.2016.03.013

44. Parra, D., Reyes-Lopez, F.E., Tort, L., 2015. Mucosal Immunity and B Cells in Teleosts: Effect of Vaccination and Stress. Front. Immunol. 6. 10.3389/fimmu.2015.00354

45. Ramadoss, N.S., Robinson, W.H., 2020. Characterizing the BCR repertoire in immune-mediated diseases. Nat. Rev. Rheumatol. 16, 7–8. 10.1038/s41584-019-0339-y

46. Rhee, K.-J., Sethupathi, P., Driks, A., Lanning, D.K., Knight, K.L., 2004. Role of commensal bacteria in development of gut-associated lymphoid tissues and preimmune antibody repertoire. J. Immunol. 172, 1118–1124. 10.4049/JIMMUNOL.172.2.1118

47. Robledo, D., Ronza, P., Harrison, P.W., Losada, A., Bermúdez, R., Pardo, B.G., José Redondo, M., Sitjà-Bobadilla, A., Quiroga, M., Martínez, P., 2014. RNA-seq analysis reveals significant transcriptome changes in turbot (Scophthalmus maximus) suffering severe enteromyxosis. BMC Genomics 15, 1149. 10.1186/1471-2164-15-1149

48. Rodriguez, O.L., Safonova, Y., Silver, C.A., Shields, K., Gibson, W.S., Kos, J.T., Tieri, D., Ke, H., Jackson, K.J.L., Boyd, S.D., Smith, M.L., Marasco, W.A., Watson, C.T., 2023. Genetic variation in the immunoglobulin heavy chain locus shapes the human antibody repertoire. Nat. Commun. 14. 10.1038/s41467-023-40070-x

49. Ronza, P., Robledo, D., Bermúdez, R., Losada, A.P., Pardo, B.G., Sitjà-Bobadilla, A., Quiroga, M.I., Martínez, P., 2016. RNA-seq analysis of early enteromyxosis in turbot (Scophthalmus maximus): new insights into parasite invasion and immune evasion strategies. Int. J. Parasitol. 46, 507–517. 10.1016/j.ijpara.2016.03.007

50. Safonova, Y., Shin, S.B., Kramer, L., Reecy, J., Watson, C.T., Smith, T.P.L., Pevzner, P.A., 2022. Variations in antibody repertoires correlate with vaccine responses. Genome Res. 32, 791–804. 10.1101/gr.276027.121

51. Sheng, X., Chai, B., Wang, Z., Tang, X., Xing, J., Zhan, W., 2019. Polymeric immunoglobulin receptor and mucosal IgM responses elicited by immersion and injection vaccination with inactivated Vibrio anguillarum in flounder (Paralichthys olivaceus). Aquaculture 505, 1–11. 10.1016/j.aquaculture.2019.02.045

52. Tang, X., Du, Y., Sheng, X., Xing, J., Zhan, W., 2018. Molecular cloning and expression analyses of immunoglobulin tau heavy chain (IgT) in turbot, Scophthalmus maximus. Vet. Immunol. Immunopathol. 203, 1–12. 10.1016/j.vetimm.2018.07.011

53. Turchaninova, M.A., Davydov, A., Britanova, O. V., Shugay, M., Bikos, V., Egorov, E.S., Kirgizova, V.I., Merzlyak, E.M., Staroverov, D.B., Bolotin, D.A., Mamedov, I.Z., Izraelson, M., Logacheva, M.D., Kladova, O., Plevova, K., Pospisilova, S., Chudakov, D.M., 2016. High-quality full-length immunoglobulin profiling with unique molecular barcoding. Nat. Protoc. 11, 1599–1616. 10.1038/nprot.2016.093

54. Vander Heiden, J.A., Yaari, G., Uduman, M., Stern, J.N.H., O’connor, K.C., Hafler, D.A., Vigneault, F., Kleinstein, S.H., 2014. pRESTO: a toolkit for processing high-throughput sequencing raw reads of lymphocyte receptor repertoires. Bioinformatics 30, 1930–1932. 10.1093/bioinformatics/btu138

55. Watson, C.T., Kos, J.T., Gibson, W.S., Newman, L., Deikus, G., Busse, C.E., Smith, M.L., Jackson, K.J.L., Collins, A.M., 2019. A comparison of immunoglobulin IGHV, IGHD and IGHJ genes in wild-derived and classical inbred mouse strains. Immunol. Cell Biol. 97, 888–901. 10.1111/imcb.12288

56. Xu, Z., Parra, D., Gomez, D., Salinas, I., Zhang, Y.-A., von Gersdorff Jorgensen, L., Heinecke, R.D., Buchmann, K., LaPatra, S., Sunyer, J.O., 2013. Teleost skin, an ancient mucosal surface that elicits gut-like immune responses. Proc. Natl. Acad. Sci. 110, 13097–13102. 10.1073/pnas.1304319110

57. Xu, Z., Takizawa, F., Parra, D., Gómez, D., von Gersdorff Jørgensen, L., LaPatra, S.E., Sunyer, J.O., 2016. Mucosal immunoglobulins at respiratory surfaces mark an ancient association that predates the emergence of tetrapods. Nat. Commun. 7, 10728. 10.1038/ncomms10728

58. Yasuike, M., de Boer, J., von Schalburg, K.R., Cooper, G.A., McKinnel, L., Messmer, A., So, S., Davidson, W.S., Koop, B.F., 2010. Evolution of duplicated IgH loci in Atlantic salmon, Salmo salar. BMC Genomics 11, 486. 10.1186/1471-2164-11-486

59. Ye, J., Ma, N., Madden, T.L., Ostell, J.M., 2013. IgBLAST: an immunoglobulin variable domain sequence analysis tool. Nucleic Acids Res. 41. 10.1093/nar/gkt382

60. Zhang, X.-T., Yu, Y.-Y., Xu, H.-Y., Huang, Z.-Y., Liu, X., Cao, J.-F., Meng, K.-F., Wu, Z.-B., Han, G.-K., Zhan, M.-T., Ding, L.-G., Kong, W.-G., Li, N., Takizawa, F., Sunyer, J.O., Xu, Z., 2021. Prevailing Role of Mucosal Igs and B Cells in Teleost Skin Immune Responses to Bacterial Infection. J. Immunol. 206, 1088–1101. 10.4049/jimmunol.2001097

61. Zhang, Y.-A., Salinas, I., Li, J., Parra, D., Bjork, S., Xu, Z., LaPatra, S.E., Bartholomew, J., Sunyer, J.O., 2010. IgT, a primitive immunoglobulin class specialized in mucosal immunity. Nat. Immunol. 11, 827–835. 10.1038/ni.1913

62. Zhu, Y., Watson, C., Safonova, Y., Pennell, M., Bankevich, A., 2025. CloseRead: a tool for assessing assembly errors in immunoglobulin loci applied to vertebrate long-read genome assemblies. Genome Biol. 26. 10.1186/s13059-025-03594-7

